# Brain-based predictions of psychiatric illness-linked behaviors across the sexes

**DOI:** 10.1101/2022.12.18.520947

**Authors:** Elvisha Dhamala, Leon Qi Rong Ooi, Jianzhong Chen, Jocelyn A. Ricard, Emily Berkeley, Sidhant Chopra, Yueyue Qu, Connor Lawhead, B.T. Thomas Yeo, Avram J. Holmes

## Abstract

**Background:** Individual differences in functional brain connectivity can be used to predict both the presence of psychiatric illness and variability in associated behaviors. However, despite evidence for sex differences in functional network connectivity and in the prevalence, presentation, and trajectory of psychiatric illnesses, the extent to which disorder-relevant aspects of network connectivity are shared or unique across the sexes remains to be determined.

**Methods:** In this work, we used predictive modelling approaches to evaluate whether shared or unique functional connectivity correlates underlie the expression of psychiatric illness-linked behaviors in males and females in data from the Adolescent Brain Cognitive Development study (n=5260; 2571 females).

**Results:** We demonstrate that functional connectivity profiles predict individual differences in externalizing behaviors in males and females, but only predict internalizing behaviors in females. Furthermore, models trained to predict externalizing behaviors in males generalize to predict internalizing behaviors in females, and models trained to predict internalizing behaviors in females generalize to predict externalizing behaviors in males. Finally, the neurobiological correlates of many behaviors are largely shared within and across sexes: functional connections within and between heteromodal association networks including default, limbic, control, and dorsal attention networks are associated with internalizing and externalizing behaviors as well as attentional deficits.

**Conclusions:** Taken together, these findings suggest that shared neurobiological patterns may manifest as distinct behaviors across the sexes. These results highlight the need to consider factors beyond just neurobiology in the diagnosis and treatment of psychiatric illnesses.

## Introduction

A primary aim of research in psychiatry is to establish the neurobiological correlates of illness-relevant behaviors, facilitating illness prediction, diagnosis, and treatment. Critical to this goal is the consideration of associated demographic characteristics, for instance underlying sex differences. Females are more likely to be diagnosed with affective and anxiety disorders, while males are more likely to meet diagnostic criteria for antisocial and substance use disorders(1–3). Relatedly, across cultures, females are more likely to express internalizing behaviors directed at one-self (i.e., loneliness, unexplained physical symptoms) while males are more likely to exhibit externalizing behaviors directed at others or the environment (i.e., aggression, hyperactivity)(3, 4). These differences emerge across childhood, become more evident during adolescence, and persist throughout the lifespan(2). While sex differences in the prevalence and expression of psychiatric illnesses have been extensively studied at the population-level(5), the underlying neurobiological correlates are not yet fully understood. Genetics, hormones, immunology, neurobiology, environment, and a host of psychosocial factors all likely contribute to expressed behaviors and these contributions may vary across disorders and throughout the lifespan(2). One possibility is that these factors uniquely contribute to distinct biological underpinnings and associated behavioral expression patterns across the sexes. An alternative, but not mutually exclusive possibility, is that shared biological features may link to dissociable behaviors across the sexes. A thorough understanding of the sex differences that exist in the neurobiological correlates of psychiatric illness-relevant behaviors will facilitate the development and implementation of sex-specific and personalized preventative interventions, diagnostic procedures, and therapeutic treatments.

Functional magnetic resonance imaging is a non-invasive neuroimaging technique that can be used to estimate regional neural activation, as inferred though the detection of changes in blood oxygenation levels. Temporal dependency patterns between these signals can subsequently be used to quantify the functional coupling (or connectivity) between pairs of brain regions. Functional connectivity profiles exhibit sex differences throughout the lifespan(6–11). Females have greater within-network connectivity while males have greater between-network connectivity(8). These differences are in part modulated by genetics(12) and hormonal fluctuations(13–16), but also likely reflect other biological, social, and environmental influences. Prior analyses have found that functional connections within and between heteromodal networks, and particularly the default and frontoparietal control networks, are largely driving these differences(9, 11). Intriguingly, functional disruptions within and between the default and control networks, along with the salience network, are also implicated in a wide range of psychiatric phenotypes(17). While sex differences in functional connectivity have been established, it is not yet known whether there are sex differences in the associations between functional connectivity and psychiatric illness-linked behaviors.

Over the last decade, data-driven predictive modeling approaches have become increasingly used to study brain-behavior relationships in healthy and clinical populations(18). These approaches can be used to not only generate individual-level clinically informative predictions of diagnosis, symptom profile, and treatment response but also to identify the underlying neurobiology that is associated with distinct psychiatric illnesses and behaviors(18). Through these approaches, functional connectivity can be used to predict individual differences in cognition, personality, as well as psychiatric and behavioral problems(19–24). These models have been used to establish the neurobiological correlates of attention(25, 26), memory(27), anxiety(28), depression(29), psychosis(30), and substance abuse(31, 32). When developing predictive models, it is crucial to ensure that they are not only accurate within circumscribed groups but that they can also generalize to other populations. Prior work indicates that predictions of cognitive and personality traits can fail to generalize across sexes(20, 33–35). To circumvent these issues–and given the known sex differences in psychiatric illnesses and behaviors–the use of sex-specific prediction models may yield more accurate and generalizable predictions and provide insight into underlying sex differences in the neurobiological correlates of psychiatric illnesses. Moreover, the examination of these brain-behavior relationships in children can reveal whether sex differences emerge prior to adolescence when many of the differences in psychiatric illness risk and presentation begin to become more evident.

Here, we sought to identify whether shared or unique neurobiological correlates underlie the expression of distinct psychiatric behaviors across the sexes during childhood. To directly address this open question, we quantified the functional connectivity correlates of 17 distinct psychiatric illness-relevant behaviors in typically developing children from the Adolescent Brain Cognitive Development (ABCD) dataset. First, by examining differences in predictive accuracy across sexes and behaviors, we demonstrate that externalizing behaviors can be accurately predicted in males and females, but internalizing behaviors can only be successfully predicted in females. Next, evaluating the generalizability of predictive models across sexes and behaviors, we determine that predictive models generalize within internalizing and externalizing domains within sexes, but only generalize across domains between sexes. More specifically, models trained to predict externalizing behaviors in either sex generalize to predict other related behaviors in both sexes. However, models trained to predict externalizing behaviors in males also generalize to predict internalizing behaviors in females, and models trained to predict internalizing behaviors in females generalize to predict externalizing behaviors in males. Finally, investigating the network correlates of these behaviors, we reveal that functional connectivity within and between shared heteromodal association networks are associated with internalizing and externalizing behaviors, as well as attention deficits, and these brain-behavior correlates are shared across the sexes. Collectively, these results suggest that shared aspects of neurobiology may underlie distinct behaviors across the sexes. Based on these findings, we encourage clinicians and researchers to consider sex when developing predictive models to facilitate diagnosis, treatment, and research of psychiatric illnesses.

## Methods

An overview of our experimental workflow is shown in Figure 1.

**Figure 1:**
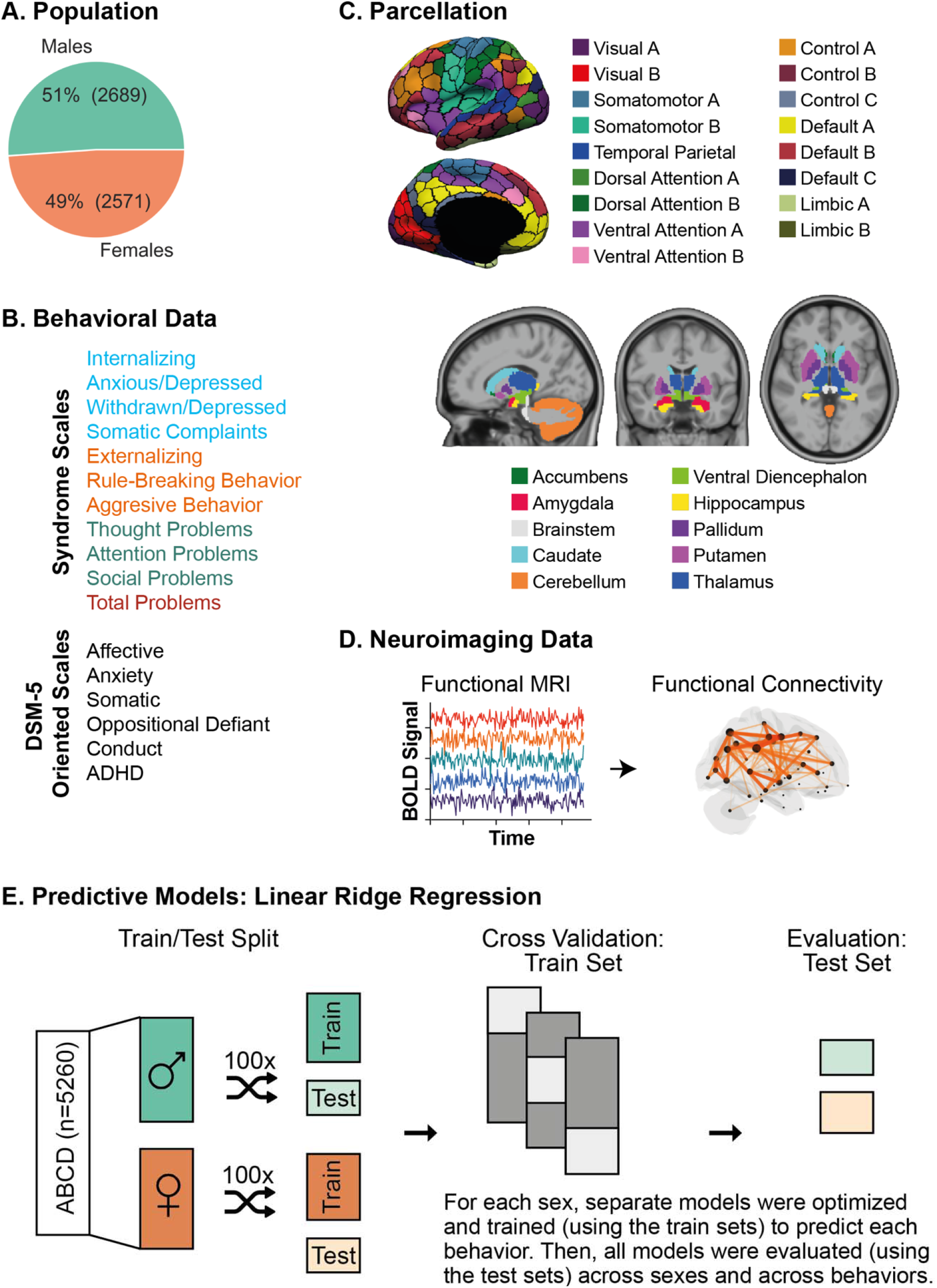
Experimental Workflow. (A) Population: We included 5260 typically developing children (9-10 years old) from the Adolescent Brain Cognitive Development (ABCD) dataset, including 2689 males (51%) and 2571 females (49%). (B) Behavioral Data: We included 17 behavioral scores from the Child Behavior Checklist which includes syndrome scales and DSM-5 oriented scales. Syndrome scales included measures of composite and individual internalizing behaviors (shown in blue), composite and individual externalizing behaviors (shown in orange), other problems (shown in green), and a summary score of total problems (red). DSM-5 Oriented Scales included scores relating to affective, anxiety, somatic, oppositional defiant, conduct, and attention deficit/hyperactivity (ADHD) disorders. (C) Parcellation: We used the Schaefer cortical parcellation of 400 regions, and each region was assigned to one of 17 large-scale cortical networks. Image reproduced under a CC BY 4.0 license: https://doi.org/10.6084/m9.figshare.10062482.v1. We also included 19 subcortical regions in our analyses, which were assigned to a subcortical network. Image reproduced under a CC BY 4.0 license: https://doi.org/10.6084/m9.figshare.10063016.v1. (D) Neuroimaging Data: For each subject, we extracted their functional MRI time series data for the 400 cortical parcels and 19 subcortical parcels. Pairwise correlation was computed for all pairs of time series to obtain the estimated functional connectivity. (E) Predictive Models: Linear ridge regression models were trained to predict individual behavioral scores based on the upper triangular functional connectivity matrix in a sex-specific manner. Data were split into training and test sets. For each training set, a separate model was optimized and trained to predict each behavior. Once optimized and trained, models were evaluated across sexes and across behaviors using the test sets.

### Dataset

We included children from the Adolescent Brain Cognitive Development (ABCD) release(36). The ABCD dataset is a large community-based sample of children and adolescents who were assessed on a comprehensive set of neuroimaging, behavioral, developmental, and psychiatric batteries. After pre-processing quality control of imaging data, as described in(22, 37), we filtered participants based on availability of functional MRI scans and behavioral scores of interest. As recommended by the ABCD consortium, we excluded individuals who were scanned using Philips scanners due to incorrect preprocessing (https://github.com/ABCD-STUDY/fMRI-cleanup). Finally, we excluded siblings to prevent unintended biases due to inherent heritability in neurobiological and/or behavioral measures. Our final ABCD sample (Figure 1A) comprised 5260 children (2689 males, 2571 females; 9-10 years old).

### Behavioral Data

The Child Behavior Checklist is a widely used clinical scale for identifying problematic behaviors in children and adolescents(38), and includes eight empirically-based syndrome scales: Anxious/Depressed, Withdrawn/Depressed, Somatic Complaints, Social Problems, Thought Problems, Attention Problems Rule-Breaking Behavior, and Aggressive Behavior. These scores are further summarized into Internalizing, Externalizing, and Total Problems. The Internalizing domain summarizes Anxious/Depressed, Withdrawn/Depressed, and Somatic Complaints. The Externalizing domains summarizes Rule-Breaking and Aggressive Behaviors. Finally, the Total Problems score is based on responses to all of the eight syndrome scales. The CBCL also includes six Diagnostic and Statistical Manual of Mental Disorders (DSM)-oriented scales consistent with DSM-5 categories: Affective (Depressive), Anxiety, Somatic, Oppositional Defiant, Conduct, and Attention Deficit/Hyperactivity (ADHD) Disorders. In these analyses, we included all eight syndrome scales, three summary scores, and six DSM-5 oriented scales for a total of 17 behavioral scores for each participant (Figure 1B). We used non-parametric Mann-Whitney U rank test to evaluate sex differences in each of the behavioral scores. All p-values were corrected for multiple comparisons using the Benjamini-Hochberg False Discovery Rate (q=0.05) procedure(39). We also computed non-parametric correlations between the behavioral scores for each sex to evaluate any underlying relationships that may exist between the behavioral scores and influence subsequent analyses.

### Image Acquisition and Processing

MR images were acquired across 21 sites in the United States using harmonized protocols for GE and Siemens scanners. The functional MRI data were preprocessed as previously described(22, 40) using a field-standard approach. Once processed, we extracted regional functional MRI time series for 400 cortical(41) and 19 subcortical(42) parcels (Figure 1C). Full correlations were then computed between those time series yielding a 419×419 pairwise regional functional connectivity matrix for each participant (Figure 1D).

### Predictive Modelling

Linear regression models and deep learning algorithms achieve comparable accuracies for brain-based behavioral predictions(23), but linear models avoid overfitting, are more interpretable, and are less computationally expensive(18). The predictive models used here rely on a similar framework as those previously described(19, 20, 43) to perform novel analyses addressing cross-behavioral model generalization within and across the sexes in the context of psychiatric illness-linked behaviors. We used linear ridge regression models to predict each behavioral score based on functional connectivity data (Figure 1E). For each sex, we split the data into 100 distinct train and test sets (at approximately a 2:1 ratio) without replacement. Imaging site was considered when splitting the data such that we placed all participants from a given site either in the train or test set but not split across the two. Within each train set, we optimized the regularization parameter using three-fold cross-validation while similarly accounting for imaging site as in the initial train-test split. Once optimized, we evaluated models on the corresponding test set. We repeated this process for each of 100 distinct train-test splits to obtain a distribution of prediction accuracy. Prediction accuracy is defined as the correlation between the true and predicted behavioral scores in the test set for each split. We computed average accuracy by taking the mean across the 100 distinct train-test splits. Once models were trained and tested within sexes and behaviors, we evaluated model generalizability across both sexes and all 17 behavioral scores. Model generalizability is defined as the accuracy obtained when a given model is evaluated on a population (i.e., sex) and/or behavioral score that is unique from the population/behavioral score that the model was trained on. This is distinct from model accuracy which is defined as the prediction accuracy obtained when evaluating the model on the same populations (i.e., sex) and behavioral score (using a hold-out test set) that it was trained on.

### Model Significance

We evaluated whether models performed better than chance levels using null distributions of performance as previously described(44). For each set of predictive models, a corresponding set of null models was generated as follows: the behavioral score was randomly permuted 1000 times, and each permutation was used to train and test a null model using a randomly selected regularization parameter from the set of selected parameters from the original model. Prediction accuracy from each of the null models was then compared to the average accuracy from the corresponding distribution of model accuracies and model generalizabilities from the original (true) models. The p-value for each model’s significance is defined as the proportion of null models with prediction accuracies greater than or equal to corresponding average accuracy from the original (true) distribution. All p-values were corrected for multiple comparisons across all measures of model accuracy and generalizability (i.e., 17 train behaviors x 2 train sexes x 17 test behaviors x 2 test sexes = 1156 comparisons) using the Benjamini-Hochberg False Discovery Rate (q=0.05) procedure(39).

### Feature Weights

We used the Haufe transformation(45) to transform feature weights obtained from the linear ridge regression models to increase their interpretability and reliability(22, 40, 46). For each train split, we used feature weights obtained from the model, *W*, the covariance of the input data (functional connectivity), *Σ*_*x*_, and the covariance of the output data (behavioral score), *Σ*_*y*_, to compute the Haufe-transformed feature weights, *A*, as follows:

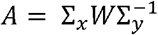

We then averaged these Haufe-transformed feature weights across the 100 splits to obtain a mean feature importance value. We computed full correlations between mean feature importance obtained from the different models to evaluate whether they relied on shared or unique features to predict the behavioral scores. For all models, we also summarized pairwise regional feature importance at a network-level to support interpretability as previously described(20). Briefly, cortical parcels were assigned to one of 17 networks from the Yeo 17-network parcellation(47), and subcortical, brainstem, and cerebellar parcels were assigned to a single subcortical network for convenience. Regional pairwise positive and negative feature weights were separately averaged to yield network-level estimates of positive and negative associations between functional connectivity and behavioral scores.

### Data and Code Availability

All ABCD data used are openly available and can be accessed directly via the NIMH Data Archive (NDA). The processed FC matrices used here were generated as part of(40) and will be uploaded to the NDA [link to be updated]. All code used to generate the results are available on GitHub [link to be updated].

## Results

### Males and females exhibit largely overlapping behaviors

The distributions of all behavioral scores included in this study are plotted for each sex in Figure 2A. While males and females exhibited largely overlapping distributions of behavioral scores, there were statistically significant (corrected p<0.01) sex differences in somatic complaints, externalizing, rule-breaking behavior, aggressive behavior, thought problems, attention problems, total problems as measured by the syndrome scales, as well as affective, somatic, oppositional defiant, conduct, and ADHD from the DSM-5 oriented scales. Males reported greater externalizing, rule-breaking behavior, aggressive behavior, thought problems, attention problems, and total problems as per the syndrome scales and greater affective, oppositional defiant, conduct, and ADHD as per the DSM-5 oriented scales. Females reported greater somatic complaints as per the syndrome scale and greater somatic problems as per the DSM-5 oriented scale.

**Figure 2:**
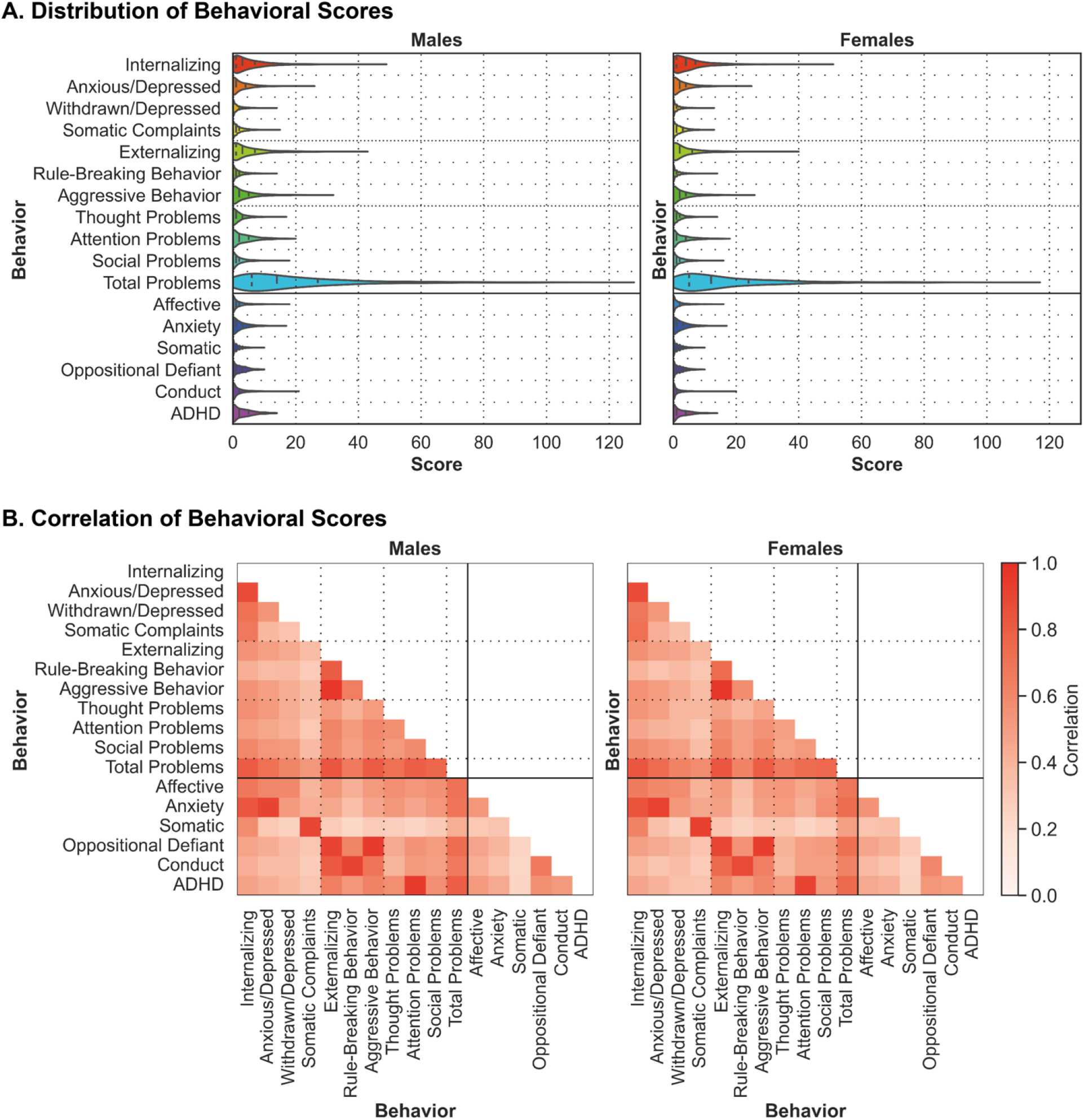
Males and females exhibit similar behavioral trends. (A) Violin plots display the distribution of all behavioral scores for males (left) and females (right). The shape of the violin plots indicates the entire distribution of values, dashed lines indicate the median, and dotted lines indicate the interquartile range. (B) The 2D grids display the correlation coefficient for each pair of behavioral scores for males (left) and females (right). ADHD – Attention deficit/hyperactivity disorder.

Within each sex, behavioral scores were strongly correlated within behavioral domains. Correlations between internalizing scores (internalizing, anxious/depressed, withdrawn/depressed, somatic complaints, affective, anxiety, somatic) ranged from 0.27 to 0.91 in males, and between 0.28 and 0.92 in females. Correlations between externalizing scores (externalizing, rule-breaking behavior, aggressive behavior, oppositional defiant, conduct) ranged between 0.53 and 0.96 in males, and between 0.51 and 0.96 in females. Correlations between attentional scores (thought problems, attention problems, social problems, ADHD) ranged between 0.52 and 0.94 in males, and between 0.48 and 0.91 in females. Meanwhile, correlations across behavioral domains were generally numerically weaker. Correlations between internalizing and externalizing scores ranged between 0.23 and 0.55 in males, and between 0.25 and 0.52 in females. Similar ranges of correlations were observed between internalizing and attentional scores, as well as externalizing and attentional scores.

Here, we replicate prior findings demonstrating sex differences in the prevalence of behaviors associated with an increased risk for illness onset and provide evidence suggesting that these differences may emerge prior to adolescence. These findings also suggest that predictive models may be more likely to generalize within sexes rather than across sexes. Additionally, we observe similar relationships between psychiatric illness-linked behaviors in males and females. These observed relationships suggest that models may be more likely to generalize within behavioral domains rather than across behavioral domains.

### Brain-based predictive models predict psychiatric illness-linked behaviors

Linear ridge regression models were trained to predict 17 psychiatric behaviors in males and females based on individual functional connectivity profiles. Once trained, model performance was evaluated in comparison to null models. Model accuracies are shown in Figure 3A.

**Figure 3:**
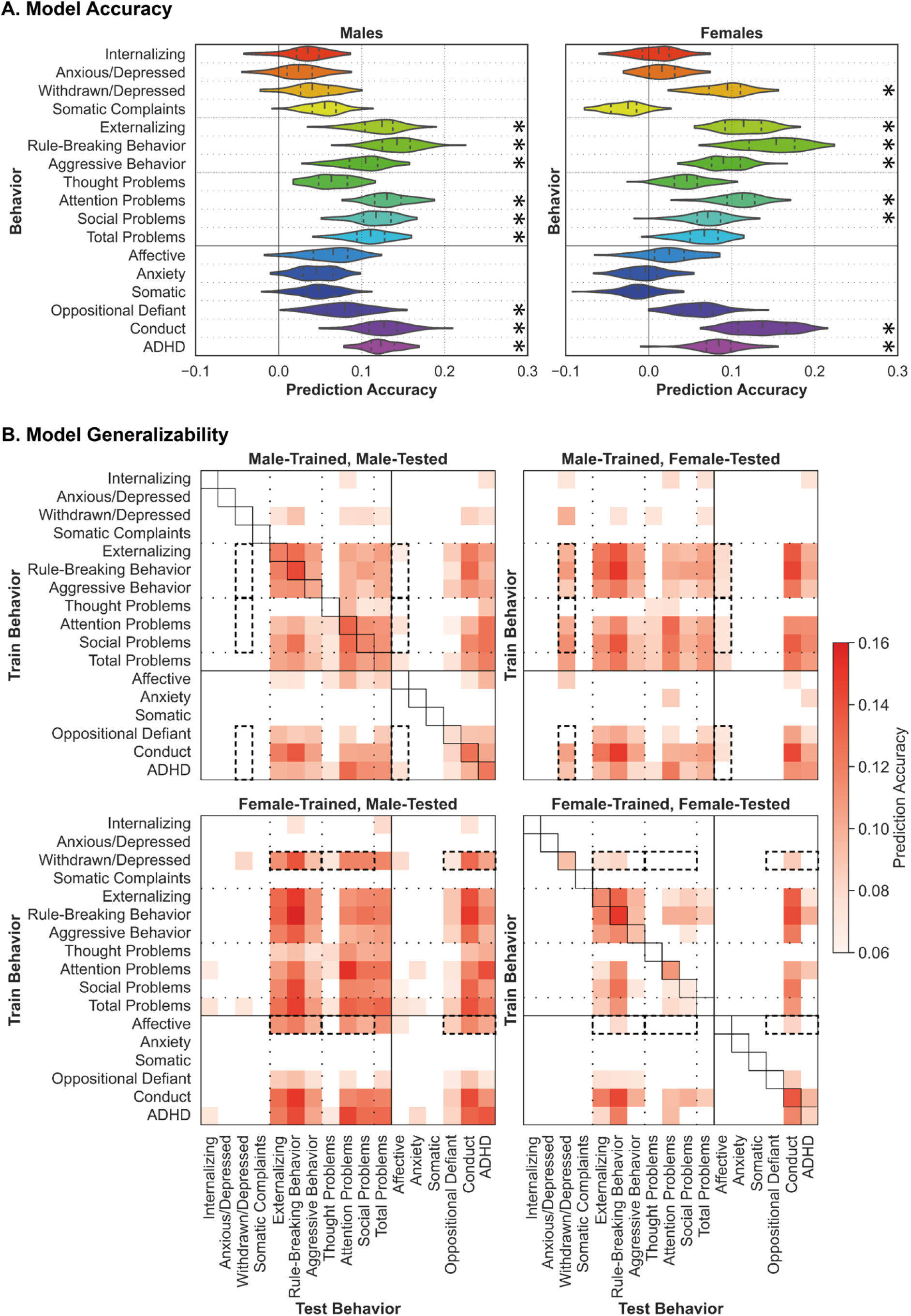
Predictive models of psychiatric illness-linked behaviors are accurate and generalizable across sexes and behaviors. (A) Model Accuracy: Model prediction accuracy (correlation coefficient between true and predicted scores) for all behaviors for males (left) and females (right). Black asterisks (*) denote that model performed significantly better than chance (corrected *p*<0.05). The shape of the violin plots indicates the entire distribution of values, dashed lines indicate the median, and dotted lines indicate the interquartile range. (B) Model Generalizability: Model generalizability across sexes and behaviors for all models. Results from models trained in males are shown at the top, and models trained in females at the bottom. Results from models tested in males are shown on the left, and models tested in females on the right. Prediction accuracy (correlation coefficient between true and predicted scores) is shown for all predictions that performed better than chance (corrected *p*<0.05) as per the color scale. For predictions that did not perform better than chance, the corresponding space is left blank. Model accuracy is shown along the diagonal for the male-trained male-tested and female-trained female-tested models (corresponding violin plots shown in Figure 3A). Dashed black boxes highlight sex differences in generalizability across behavioral domains.

In males, models successfully predicted behaviors (corrected p<0.05) within the externalizing domain (externalizing (r=0.12), rule-breaking (r=0.14), and aggressive (r=0.10) behaviors), as well as attention (r=0.13), social (r=0.12), and total (r=0.11) problems from the syndrome scales. Models also successfully predicted behaviors related to oppositional defiant (r=0.08), conduct (0.13), and attention deficit/hyperactivity (ADHD; r=0.12) disorders from the DSM-5 oriented scales in males.

In females, models successfully predicted behaviors (corrected p<0.05) within the internalizing domain (withdrawn/depressed (r=0.09)) and the externalizing domain (externalizing (r=0.11), rule-breaking (r=0.15), and aggressive (r=0.09) behaviors), as well as attention (r=0.11) and social (r=0.07) problems from the syndrome scales. Models also successfully predicted behaviors related to conduct disorders (r=0.14) and ADHD (r=0.08) from the DSM-5 oriented scales.

In our prior work, we have observed that internalizing behaviors are more difficult to predict than externalizing behaviors(22). Our results replicate these findings and further suggest that the predictability of specific behaviors may differ across the sexes.

### Brain-based predictive models of psychiatric illness-linked behaviors generalize across sexes and behaviors

Generalizability of linear ridge regression models trained in each sex to predict each of the 17 behaviors were evaluated across sexes and across behaviors. Generalizability is defined as the prediction accuracy obtained when a given model is evaluated on a population and/or behavior distinct from the population and/or behavior it was trained on. These are shown in Figure 3B.

Models trained in males (top row in Figure 3B) to predict externalizing syndromes (externalizing, rule-breaking, and aggressive behaviors), and attention, social, and total problems, as well as behaviors related to oppositional defiant and conduct disorders, and ADHD successfully generalize (corrected p<0.05) across those behaviors in males and females. These models also generalize (corrected p<0.05) to predict internalizing (withdrawn/depressed) syndromes and behaviors related to affective disorders in females, but not in males (see dashed black boxes in top row of Figure 3B). Additionally, models trained to predict internalizing syndromes (internalizing and withdrawn/depressed behaviors) and affective behaviors generalize (corrected p<0.05) to predict some externalizing syndromes as well as attention problems and behaviors related to ADHD in males and females, albeit to a weaker extent.

Models trained in females (bottom row in Figure 3B) to predict externalizing syndromes (externalizing, rule-breaking, and aggressive behaviors), attention and social problems, as well as behaviors related to conduct disorders and ADHD successfully generalize (corrected p<0.05) across those behaviors in males and females. Surprisingly, these models trained in females exhibit generally greater generalizability in males (bottom left panel in Figure 3B) than in females (bottom right panel in Figure 3B). In other words, models trained in females more accurately predict behaviors in males than in females. Moreover, models trained to predict internalizing syndromes (withdrawn/depressed) and affective behaviors generalize (corrected p<0.05) to predict externalizing syndromes (externalizing and rule-breaking behaviors), thought, attention, social, and total problems, and behaviors related to oppositional defiant and conduct disorders, and ADHD in males (see dashed boxes in bottom row of Figure 3B). Similar results are also observed when generalizing (corrected p<0.05) within females but to a lesser extent.

Taken together, these results suggest that brain-based predictive models trained in one domain can generalize to predict other related behaviors within the same domain. These models may also generalize to predict behaviors in other unrelated domains and this generalizability may be more evident across sexes rather than within sexes.

### Functional correlates of psychiatric behaviors are shared across behaviors and sexes

Pairwise regional feature weights used to predict psychiatric illness-linked behaviors were extracted from the models and Haufe-transformed. Correlations between these Haufe-transformed feature weights across both sexes and all behaviors were analyzed and are shown in Figure 4.

**Figure 4:**
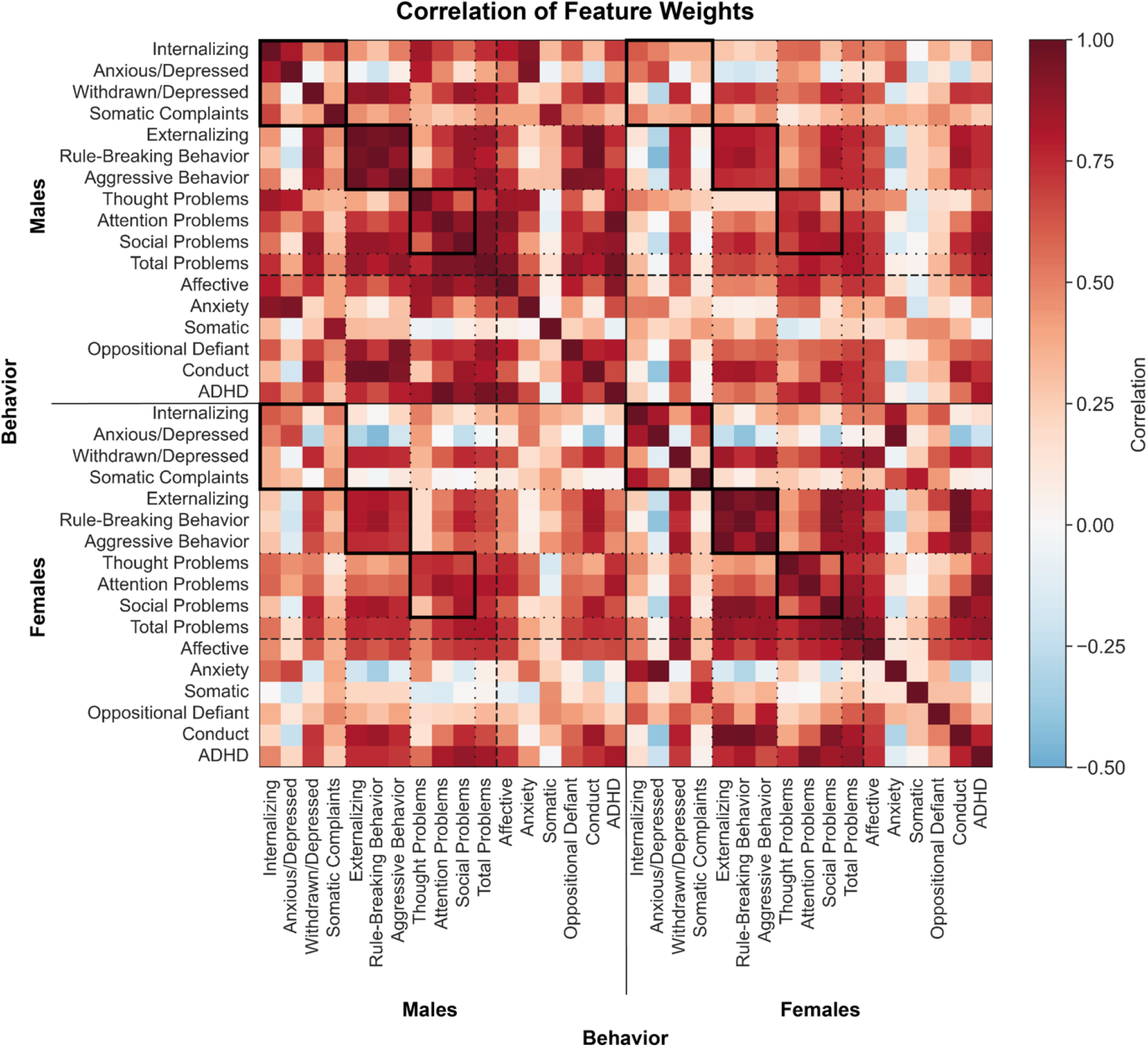
Shared functional connectivity features underlie distinct behaviors across the sexes. Correlation coefficient between Haufe-transformed pairwise regional feature weights from distinct models. Models trained in males are shown at the top and on the left, models trained in females are shown at the bottom and on the right. Warmer colors indicate a positive correlation and cooler colors indicate a negative correlation. Solid black boxes highlight correlations between feature weights within behavioral domains within and between sexes.

Feature weights are strongly correlated across behaviors and sexes, and the strongest correlations are observed within behavioral domains (see solid black boxes in Figure 4). One notable exception is the features involved in the prediction of anxious/depressed behaviors and somatic complaints, as well as anxiety and somatic diagnoses in males and females, both of which exhibit generally weak correlations with features for all other predictions including those within the internalizing domain, but strong positive correlations with each other (see rows and columns depicting correlations for Anxious/Depressed, Somatic Complaints, Anxiety, and Somatic).

In prior work, we have demonstrated that shared features predict a smaller subset of psychiatric behaviors(22). Here, we replicate those findings and demonstrate that even though males and females may exhibit behavioral differences, shared neurobiological features underlie the expression of those behaviors.

### Functional connectivity within and between different heteromodal association networks predict psychiatric illness-linked behaviors

Regional pairwise feature weights were summarized to a network-level based on the Yeo 17-network solution(47). Positive and negative feature weights were separately averaged to yield positive and negative network-level associations between functional connectivity and psychiatric behaviors. For simplicity, we show the corresponding figures for three behaviors (withdrawn/depressed, rule-breaking, attention) characteristics of the three broader psychiatric behavioral domains (internalizing, externalizing, attention) in Figures 5–7, respectively, and for all other behaviors in the supplemental materials (Figures S1-S14). Correlations between these network-level associations across the sexes are shown in Table S1.

**Figure 5:**
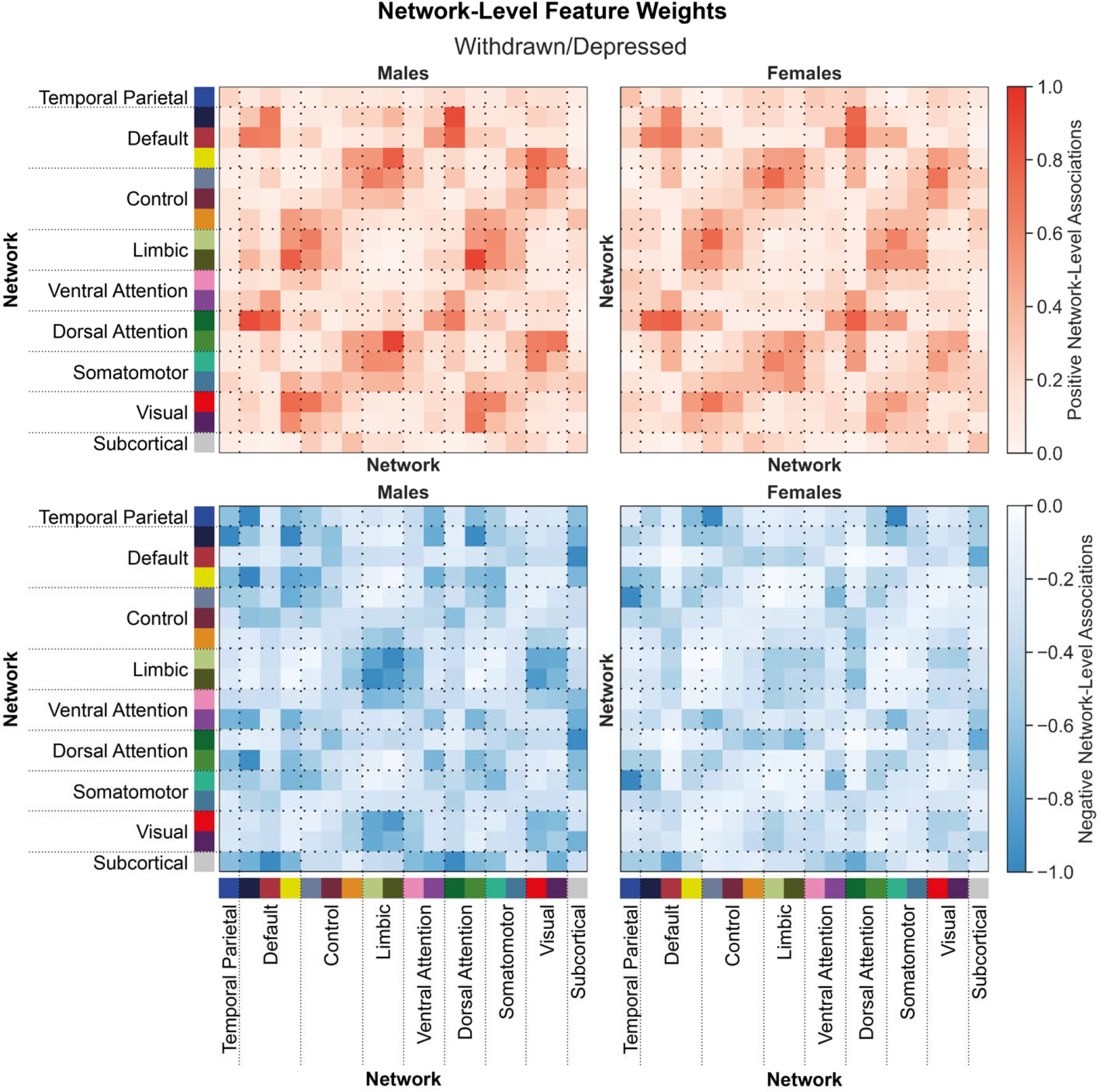
Shared network-level functional connections underlie withdrawn/depressed behaviors in males and females. Positive (top) and negative (bottom) associations between network-level functional connectivity and rule-breaking behaviors in males (left) and females (right). Regional feature weights were summarized to a network-level by assigning cortical regions to one of 17 Yeo networks, and subcortical regions to a subcortical network. Colors next to the network labels along the vertical and horizontal axes correspond to the network colors from Figure 1C. Warmer colors within the heatmap indicate a positive association and cooler colors indicate a negative association. For visualization, values within each matrix were divided by the absolute maximum value across the positive and negative matrices for each sex. Correlations between positive associations across sexes, r_positive_=0.89. Correlations between negative associations across sexes, r_negative_=0.72.

Across both sexes, functional connectivity within and between the default and dorsal attention networks are positively associated with withdrawn/depressed behaviors (Figure 5, top row). Functional connections between the limbic network and the default, control, dorsal attention, and somatomotor networks are also positively associated with withdrawn/depressed behaviors (Figure 5, top row). Finally, functional connections between the visual network and the default and dorsal attention networks are also positively associated with withdrawn/depressed behaviors (Figure 5, top row), although to a slightly weaker extent in females than in males. In males, functional connectivity within default and limbic networks, as well as between default and temporal parietal, and default and dorsal attention networks were negatively associated with withdrawn/depressed behaviors (Figure 5, bottom left). Widespread cortico-subcortical connections were also negatively associated with withdrawn/depressed behaviors in males (Figure 5, bottom left). In females, generally fewer negative associations were observed, and those observed occurred between the temporal parietal network and the control and somatomotor networks (Figure 5, bottom right). Positive and negative associations were largely shared across the sexes (r_positive_=0.89, r_negative_=0.72).

Functional connections that were associated with rule-breaking behaviors (Figure 6) were largely similar to those associated with withdrawn/depressed behaviors with a few key differences. Functional connections between the visual network and the default and dorsal attention exhibited a slightly stronger association with rule-breaking behaviors in males than in females (Figure 6, top row). Moreover, rather than widespread negative associations with cortico-subcortical connections, subcortical connections to the default and dorsal attention networks were most strongly negatively associated with rule-breaking behaviors (Figure 6, bottom row). These associations were also similar across the sexes (r_positive_=0.90 for positive, and r_negative_=0.94).

**Figure 6:**
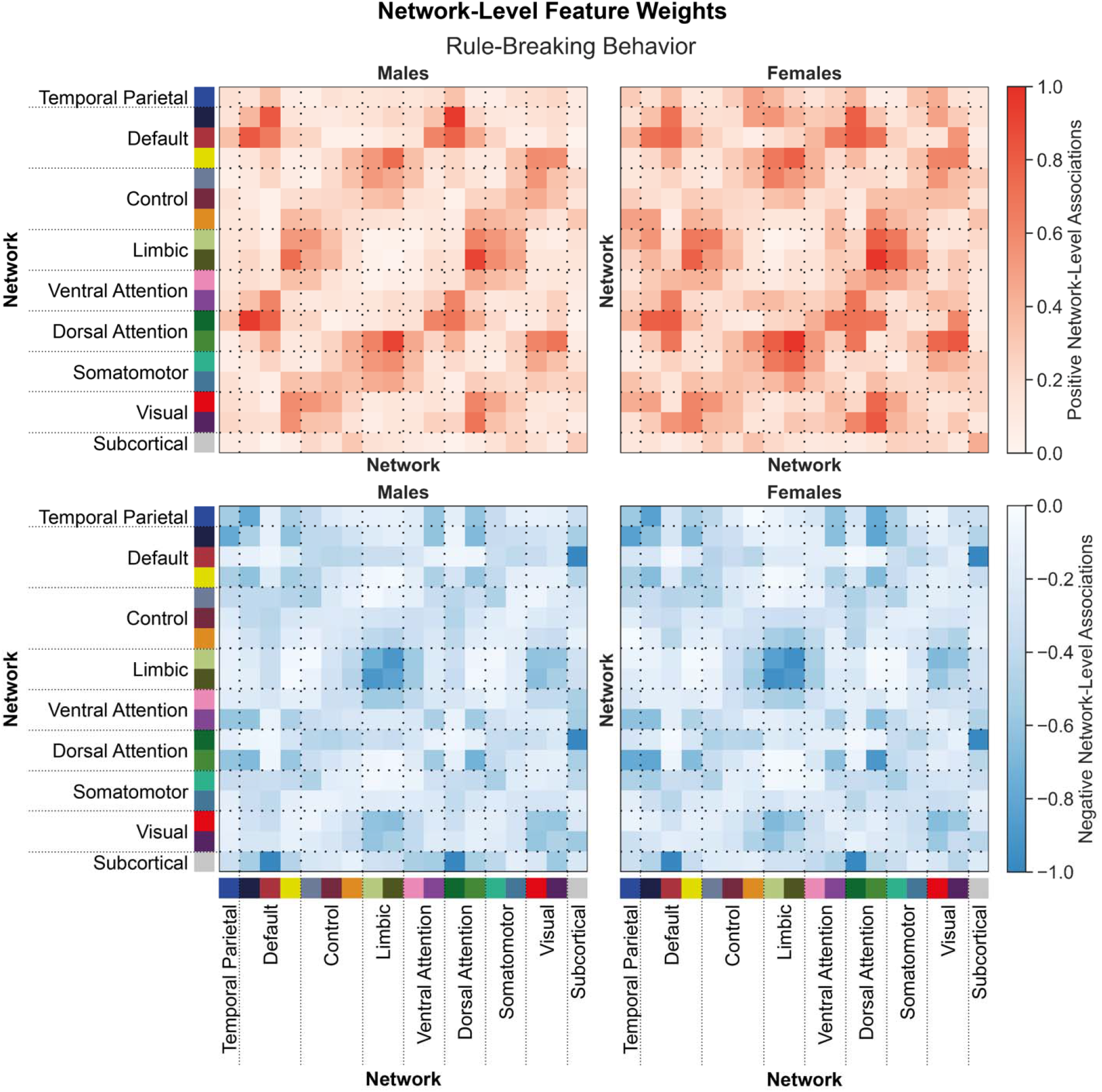
Shared network-level functional connections underlie rule-breaking behaviors in males and females. Positive (top) and negative (bottom) associations between network-level functional connectivity and rule-breaking behaviors in males (left) and females (right). Regional feature weights were summarized to a network-level by assigning cortical regions to one of 17 Yeo networks, and subcortical regions to a subcortical network. Colors next to the network labels along the vertical and horizontal axes correspond to the network colors from Figure 1C. Warmer colors within the heatmap indicate a positive association and cooler colors indicate a negative association. For visualization, values within each matrix were divided by the absolute maximum value across the positive and negative matrices for each sex. Correlations between positive associations across sexes, r_positive_=0.90. Correlations between negative associations across sexes, r_negative_=0.94.

Functional connections between the limbic network and the default, control, dorsal attention, and somatomotor networks, as well as connections between the visual network and the default, control, dorsal attention, and somatomotor networks were associated with attention problems in males and females (Figure 7, top row). We do not observe any strong negative associations between functional connectivity and attention problems (Figure 7, bottom row). Similar to the observations for the withdrawn/depressed and rule-breaking behaviors, these associations were shared across the sexes (r_positive_=0.95 for positive, r_negative_=0.95).

**Figure 7:**
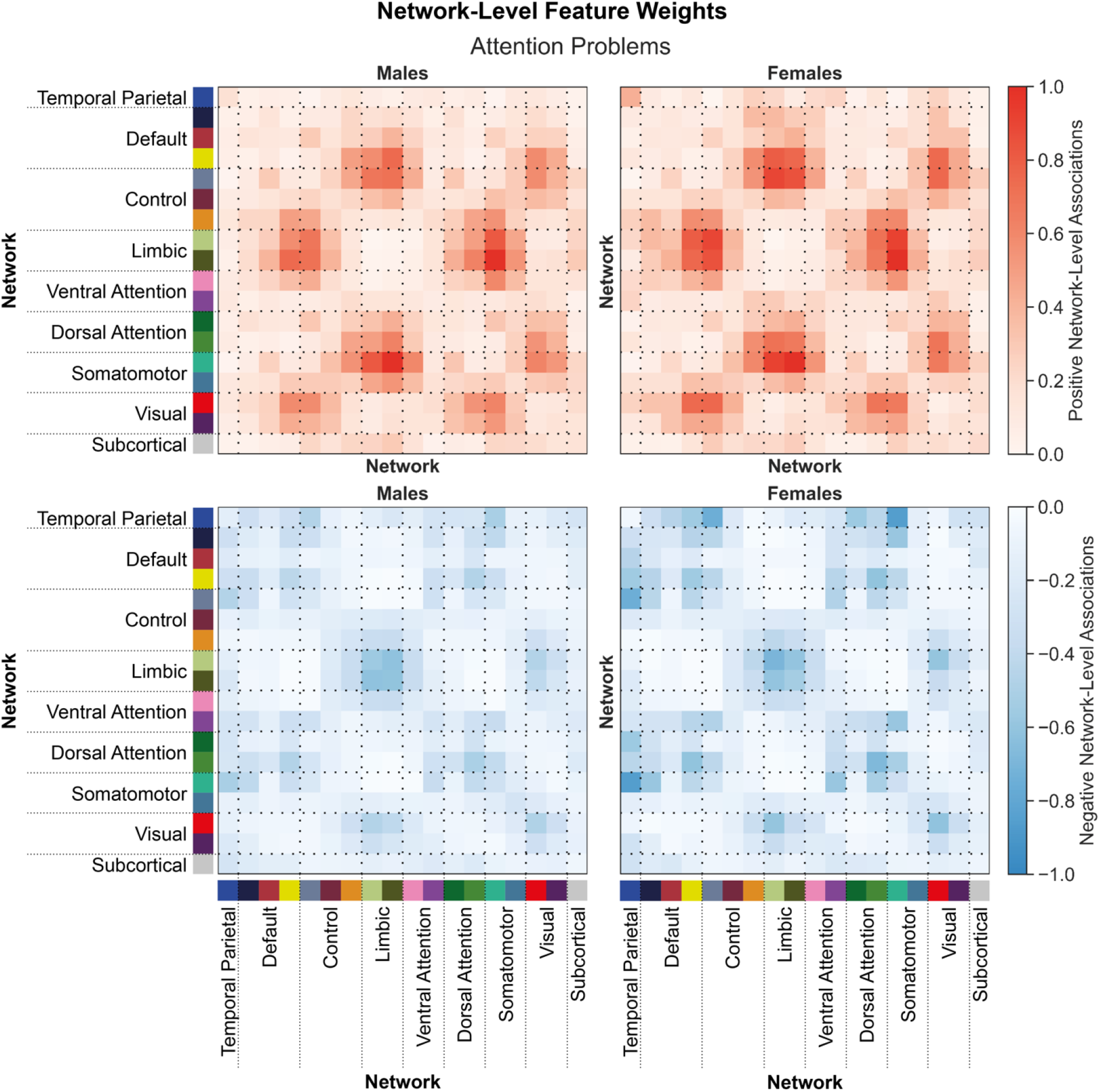
Shared network-level functional connections underlie attention problems in males and females. Positive (top) and negative (bottom) associations between network-level functional connectivity and attention problems in males (left) and females (right). Regional feature weights were summarized to a network-level by assigning cortical regions to one of 17 Yeo networks, and subcortical regions to a subcortical network. Colors next to the network labels along the vertical and horizontal axes correspond to the network colors from Figure 1C. Warmer colors within the heatmap indicate a positive association and cooler colors indicate a negative association. For visualization, values within each matrix were divided by the absolute maximum value across the positive and negative matrices for each sex. Correlations between positive associations across sexes, r_positive_=0.95. Correlations between negative associations across sexes, r_negative_=0.94.

These findings are in line with prior work demonstrating that functional connections in heteromodal association networks are largely implicated in a wide range of psychiatric illnesses(48–51). We further demonstrate that shared functional connectivity correlates underlie internalizing and externalizing behaviors across the sexes. Moreover, while there exist some similarities in the networks associated with attention problems, there are also unique network contributions observed within the attention domain. Altogether, these findings suggest that while shared neurobiological correlates are likely to be observed across psychiatric behaviors and illnesses, there are also distinct network signatures associations with different behavioral domains.

## Discussion

Brain-based predictive modeling has provided foundational insights into the neurobiological correlates of psychiatric illness(18, 52–54). While associations between functional connectivity and distinct psychiatric illnesses and behaviors have been studied extensively, prior work has not yet addressed whether those relationships are shared across the sexes. Functional connectivity profiles and the expression of psychiatric illnesses are both known to differ across males and females, but it is not clear whether these differences map onto one another. Here, we demonstrate in a large sample of 5260 children from the ABCD dataset that functional connectivity profiles predict externalizing behaviors and attention deficits in males and females, but internalizing behaviors are generally only predictable in females. Models trained to predict externalizing behaviors and attention deficits generalize across those behavioral domains within and between sexes. Moreover, models trained to predict externalizing behaviors in males can also predict internalizing behaviors in females. Likewise, models trained to predict internalizing behaviors in females can also predict externalizing behaviors in males. Across both males and females, functional connections within and between heteromodal association networks underlie the expression of internalizing and externalizing behaviors, as well as attentional deficits. Taken together, these results reveal that shared disruptions in functional connectivity can manifest as distinct psychiatric behaviors across the sexes.

Psychiatric diagnoses describe clusters of problematic behaviors that tend to overlap across diagnoses(55), lack clear discernible boundaries(55), and exhibit high rates of comorbidity(56). Consequently, it is extremely difficult to isolate disorder-specific biomarkers. To understand the neurobiological processes that underlie distinct psychiatric illnesses, several different approaches have been posited. The dimensional approach proposes that psychopathology can be described along distinct dimensions of psychiatric illness(57, 58). An individual’s vulnerability to a particular psychiatric illness can be defined by how they score across different dimensions. Similarly, the internalizing-externalizing model suggests that psychiatric illnesses are manifestations of internalizing and externalizing dimensions(59), where internalizing dimensions affect an individual’s internal state and externalizing dimensions affect an individual’s external response to the world(60). An alternative theory, the p-factor, suggests a single factor of psychopathology makes individuals broadly vulnerable to psychiatric illness and the specific illness they develop is determined by other factors(61). Regardless of how we characterize distinct psychiatric illnesses and associated behaviors, an understanding of their underlying associations with brain-based biomarkers is crucial for the development of personalized diagnostic approaches and treatment interventions. These present analyses suggest behavioral prediction models may be broadly generalizable across dimensional measures and diagnosis-based scales, increasing their clinical utility. Furthermore, by moving beyond the categorical medical model and integrating dimensional measures, we can improve our understanding of the range of psychiatric symptom profiles that may be associated with functional network connectivity.

Our prior work suggests psychiatric illnesses and associated behaviors are generally harder to predict than cognitive traits and exhibit weaker associations with neurobiological features(21, 22). Relatedly, brain-based models of internalizing behaviors and illnesses tend to achieve weaker prediction accuracies than those of externalizing behaviors and illnesses(22). The general lack of predictability of internalizing behaviors seen here and in prior work may be related to individual differences in the signal-to-noise ratio in the associations between functional connectivity and the behaviors themselves. Furthermore, the presence of significant predictions of internalizing behaviors in females, but not in males, may be underscored by the earlier development of functional networks, and especially the heteromodal association networks, in females during childhood(7, 62). The delayed development of association networks–which drive these behavioral predictions–paired with the lower levels of internalizing behaviors observed in males, could in part explain the lower observed accuracies in males.

Prior and ongoing analyses of the neurobiological correlates of psychopathology suggest functional disruptions in heteromodal association networks are implicated across dimensions and disorders: affective and psychotic illnesses as well as symptoms associated with those illnesses are related to frontoparietal control, limbic, default, and attention network connectivity(22, 48–51). In this present study, we find functional connections within and between those networks predict individual differences in psychiatric illness-linked behaviors. While connections between limbic and frontoparietal networks are associated with all behaviors analyzed, other distinct functional network signatures are associated with specific syndromes and DSM-oriented traits. These findings suggest the existence of transdiagnostic and disorder-specific functional signatures of psychiatric illnesses and illness-linked behaviors. Finally, shared genetic and environmental influences have been shown to underlie the covariant expression of negative affect, internalizing behaviors, and externalizing behaviors(63). Our results further suggest these traits may also share neurobiological influences, which may in part be driven by genetic and environmental influences on neurobiology itself.

Sex differences in neurobiology and behavior are well established(2, 5–7, 64–77). More recently, researchers have also begun to look at sex differences in brain-behavior relationships(20, 33, 34, 43, 78). To explain the underlying factors driving these differences in clinical populations, sex-based and gender-based theories have been proposed. Sex-based theories posit that sex chromosomes, brain structure, the hypothalamic-pituitary-adrenal axis, immune processes, and gonadal hormones underlie sex differences in psychiatric illnesses, while gender-based theories emphasize the contributions of parental expectations, gender socialization, gender roles, gender identities, and diagnostic biases(3). In this present study, we demonstrate functional correlates of psychiatric illness-linked behaviors are largely shared across the sexes. Furthermore, shared functional correlates are associated with the expression of internalizing and externalizing behaviors, of which, internalizing are more prevalent in females and externalizing in males. These findings suggest that differences observed in the expression of psychiatric illness-linked behaviors across the sexes are not dependent on sex-specific functional connectivity profiles, but we are not able to rule out the contributions of other sex- or gender-related factors.

The findings of this study are subject to several limitations. First, these analyses relied on a large community-based sample of children between the ages of 9 and 10. As these children undergo puberty and go through adolescence, there will likely exhibit changes in their behavioral expressions and brain biology, particularly in the heteromodal association networks(79–82). As such, the underlying brain-behavior relationships are subject to change throughout the course of adolescence. Subsequent analyses investigating brain-behavior relationships at the follow-up time points in the ABCD data could address this question. Second, since the ABCD dataset does not include information about gender identity or fluidity, this study only used information about each subject’s self-reported sex. Throughout the course of development, males and females are exposed to gender-differentiated experiences and enculturation. Given the lack of data pertaining to gender, we cannot disentangle whether the observed sex differences are driven by inherent sex differences in neurobiology and/or behavior a manifestation of gender-related differences, or a combination of the two such that innate biological differences are further exaggerated by sociocultural and environmental factors(83). Third, this study used a single dataset which was collected entirely in the United States. While the dataset was acquired using different sites (and scanners) across the country suggesting these results are somewhat generalizable, it does not represent the global extent of racial, ethnical, or cultural diversity. As such, further research is needed to address whether these results are generalizable across populations(84, 85) with known differences in the expression, diagnosis, and stigmatization of psychiatric illness-linked behaviors(86–88).

## Acknowledgements

Data used in the preparation of this article were obtained from the Adolescent Brain Cognitive Development ^SM^ (ABCD) Study (https://abcdstudy.org), held in the NIMH Data Archive (NDA). This is a multisite, longitudinal study designed to recruit more than 10,000 children aged 9-10 and follow them over 10 years into early adulthood. The ABCD Study® is supported by the National Institutes of Health and additional federal partners under award numbers U01DA041048, U01DA050989, U01DA051016, U01DA041022, U01DA051018, U01DA051037, U01DA050987, U01DA041174, U01DA041106, U01DA041117, U01DA041028, U01DA041134, U01DA050988, U01DA051039, U01DA041156, U01DA041025, U01DA041120, U01DA051038, U01DA041148, U01DA041093, U01DA041089, U24DA041123, U24DA041147. A full list of supporters is available at abcdstudy.org/federal-partners.html. A list of participating sites and a complete list of the study investigators can be found at abcdstudy.org/consortium_members/. ABCD consortium investigators designed and implemented the study and/or provided data but did not necessarily participate in the analysis or writing of this report. This manuscript reflects the views of the authors and may not reflect the opinions or views of the NIH or ABCD consortium investigators.

