## Supplemental Materials for "Brain-based predictions of psychiatric illness-linked behaviors across the sexes"

Running Title: Brain networks predict distinct behaviors across sexes

**Authors:**

Elvisha Dhamala^1-2,*^, Leon Qi Rong Ooi^3-6^**,** Jianzhong Chen^4-6^**,** Jocelyn A. Ricard^1^, Emily Berkeley^7^, Sidhant Chopra^1^, Yueyue Qu^1^, Connor Lawhead^1^, B.T. Thomas Yeo^3-6,8^**,** Avram J. Holmes^1-2,9-10,*^

**Affiliations:**

^1^ Department of Psychology, Yale University, New Haven, USA

^2^ Kavli Institute for Neuroscience, Yale University, New Haven, USA

^3^ Centre for Sleep & Cognition & Centre for Translational Magnetic Resonance Research, Yong Loo Lin School of Medicine, Singapore, National University of Singapore, Singapore, Singapore

^4^ Department of Electrical and Computer Engineering, National University of Singapore, Singapore, Singapore

^5^ N.1 Institute for Health & Institute for Digital Medicine, National University of Singapore, Singapore, Singapore

^6^ Integrative Sciences and Engineering Programme (ISEP), National University of Singapore, Singapore, Singapore

^7^ Connecticut College, New London, USA

^8^ Martinos Center for Biomedical Imaging, Massachusetts General Hospital, Charlestown, USA

^9^ Department of Psychiatry, Yale University, New Haven, USA

^10^ Wu Tsai Institute, Yale University, New Haven, USA

^*^ Corresponding authors:
Elvisha Dhamala and Avram J. Holmes

Address: 401 Sheffield Sterling Strathcona Hall, 1 Prospect Street, New Haven, CT 06511

Phone Number: 203-436-9449

**Keywords:** prediction psychiatry, neuroimaging, functional connectivity, brain-based predictions, sex differences, transdiagnostic

**Supplementary Table 1: Shared network-level features underlie psychiatric illness-linked behaviors across the sexes.**

Correlation coefficient between network-level feature weights from models trained on males and females. Correlations were computed separately for positive associations and negative associations. Corresponding network-level positive and negative associations for all behaviors for males and females are shown in Figures 5-7 and Supplementary Figures 1-14.

| **Behavior** | **Positive Associations** | **Negative Associations** |
| --- | --- | --- |
| Internalizing | 0.66 | 0.72 |
| Anxious/Depressed | 0.55 | 0.73 |
| Withdrawn/Depressed | 0.89 | 0.72 |
| Somatic Complaints | 0.78 | 0.54 |
| Externalizing | 0.87 | 0.89 |
| Rule-Breaking Behavior | 0.90 | 0.94 |
| Aggressive Behavior | 0.80 | 0.75 |
| Thought Problems | 0.91 | 0.86 |
| Attention Problems | 0.95 | 0.94 |
| Social Problems | 0.82 | 0.90 |
| Total Problems | 0.95 | 0.88 |
| Affective | 0.81 | 0.75 |
| Anxiety | 0.38 | 0.53 |
| Somatic | 0.76 | 0.56 |
| Oppositional Defiant | 0.50 | 0.27 |
| Conduct | 0.89 | 0.93 |
| ADHD | 0.91 | 0.94 |

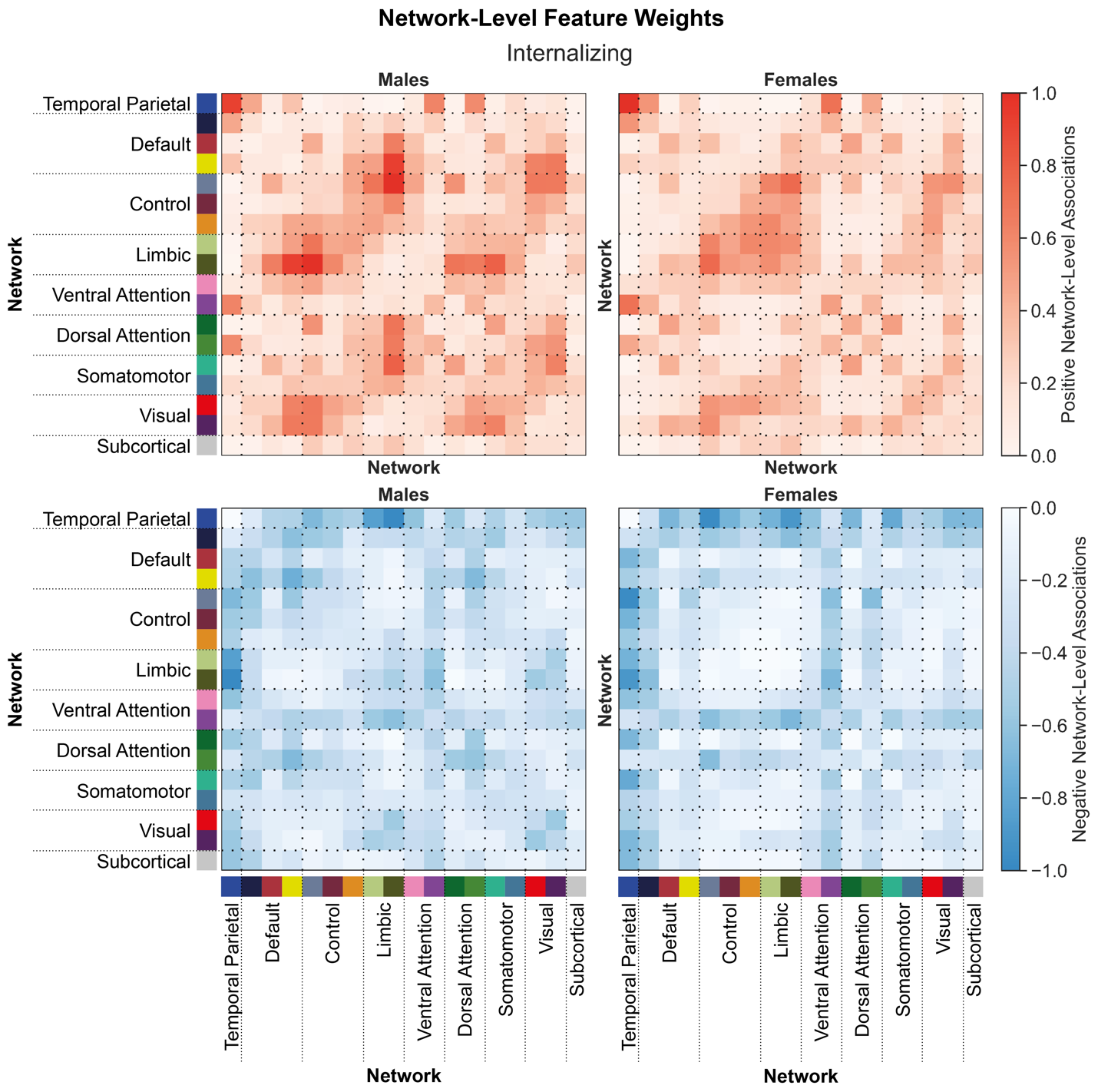
 **Supplementary Figure 1: Shared network-level functional connections underlying internalizing behaviors in males and females.**

Positive (top) and negative (bottom) associations between network-level functional connectivity and internalizing behaviors in males (left) and females (right). Regional feature weights were summarized to a network-level by assigning cortical regions to one of 17 Yeo networks, and subcortical regions to a subcortical network. Colors next to the network labels along the vertical and horizontal axes correspond to the network colors from Figure 1C. Warmer colors within the heatmap indicate a positive association and cooler colors indicate a negative association. For visualization, values within each matrix were divided by the absolute maximum value across the positive and negative matrices for each sex. Correlations between positive associations across sexes, r_positive_=0.66. Correlations between negative associations across sexes, r_negative_=0.72.

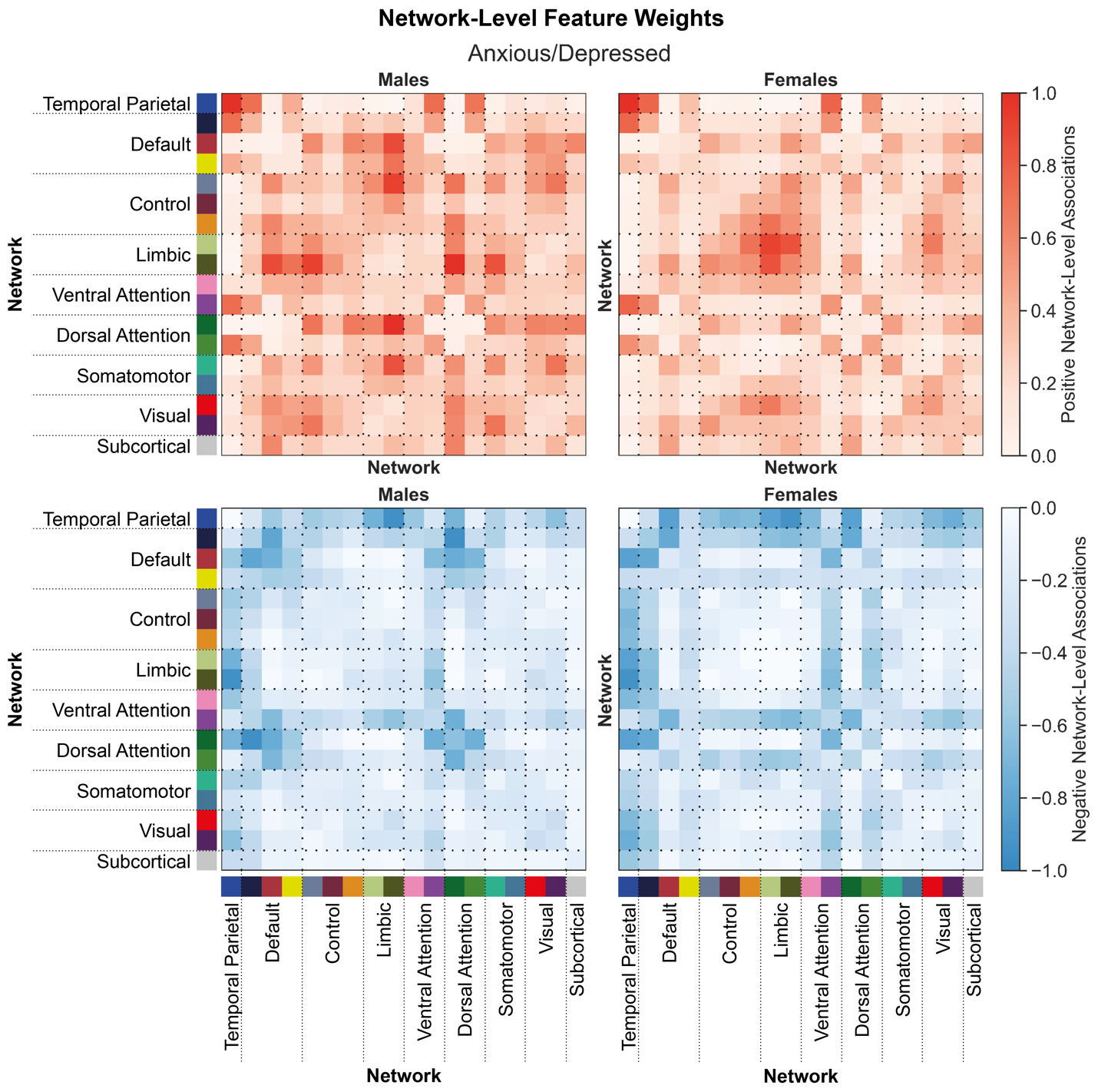
 **Supplementary Figure 2: Shared network-level functional connections underlying anxious/depressed behaviors in males and females.**

Positive (top) and negative (bottom) associations between network-level functional connectivity and anxious/depressed behaviors in males (left) and females (right). Regional feature weights were summarized to a network-level by assigning cortical regions to one of 17 Yeo networks, and subcortical regions to a subcortical network. Colors next to the network labels along the vertical and horizontal axes correspond to the network colors from Figure 1C. Warmer colors within the heatmap indicate a positive association and cooler colors indicate a negative association. For visualization, values within each matrix were divided by the absolute maximum value across the positive and negative matrices for each sex. Correlations between positive associations across sexes, r_positive_=0.55. Correlations between negative associations across sexes, r_negative_=0.73.

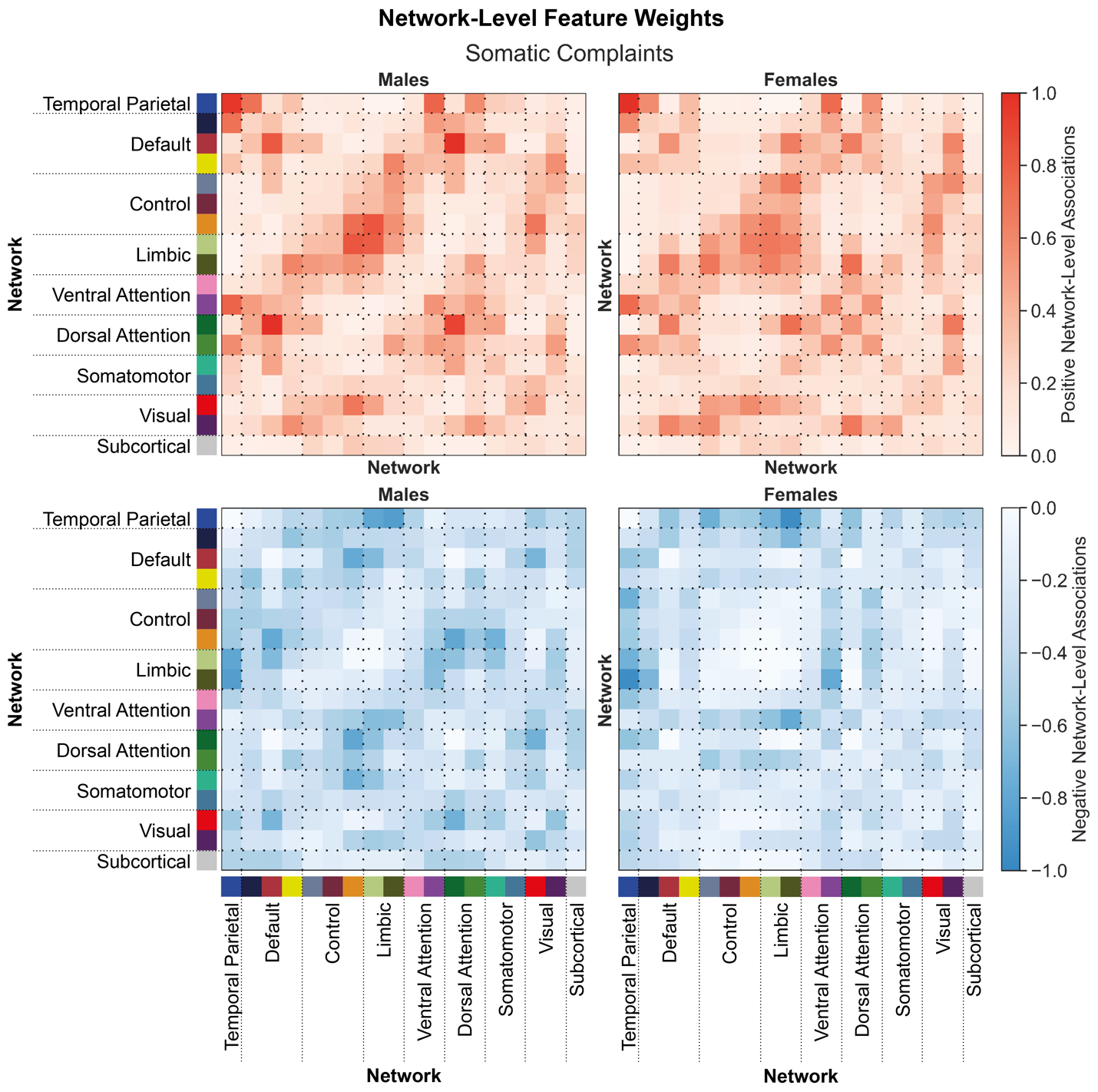
 **Supplementary Figure 3: Shared network-level functional connections underlying somatic complaints in males and females.**

Positive (top) and negative (bottom) associations between network-level functional connectivity and somatic complaints in males (left) and females (right). Regional feature weights were summarized to a network-level by assigning cortical regions to one of 17 Yeo networks, and subcortical regions to a subcortical network. Colors next to the network labels along the vertical and horizontal axes correspond to the network colors from Figure 1C. Warmer colors within the heatmap indicate a positive association and cooler colors indicate a negative association. For visualization, values within each matrix were divided by the absolute maximum value across the positive and negative matrices for each sex. Correlations between positive associations across sexes, r_positive_=0.78. Correlations between negative associations across sexes, r_negative_=0.54.

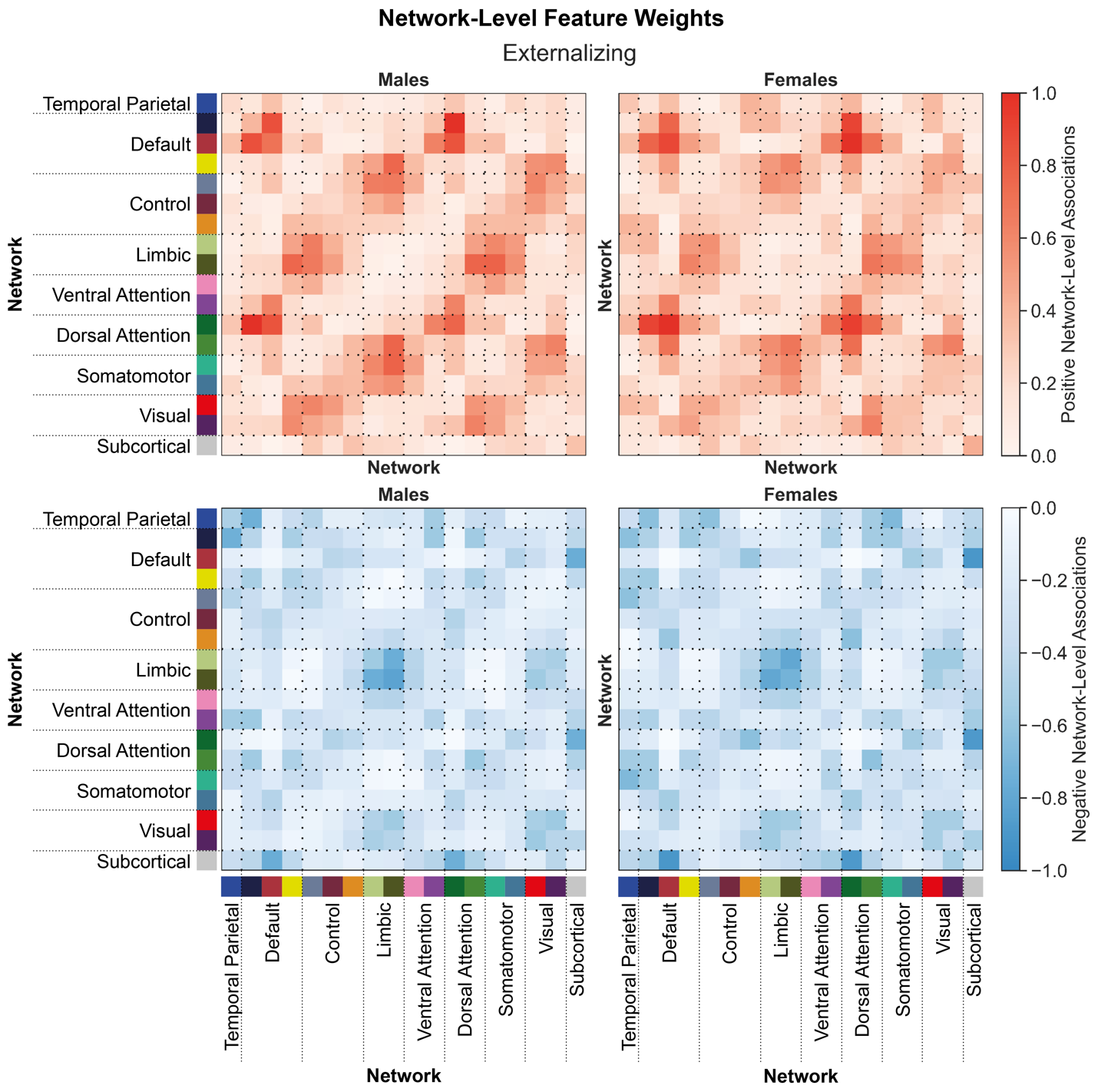
 **Supplementary Figure 4: Shared network-level functional connections underlying externalizing behaviors in males and females.**

Positive (top) and negative (bottom) associations between network-level functional connectivity and externalizing behaviors in males (left) and females (right). Regional feature weights were summarized to a network-level by assigning cortical regions to one of 17 Yeo networks, and subcortical regions to a subcortical network. Colors next to the network labels along the vertical and horizontal axes correspond to the network colors from Figure 1C. Warmer colors within the heatmap indicate a positive association and cooler colors indicate a negative association. For visualization, values within each matrix were divided by the absolute maximum value across the positive and negative matrices for each sex. Correlations between positive associations across sexes, r_positive_=0.87. Correlations between negative associations across sexes, r_negative_=0.89.

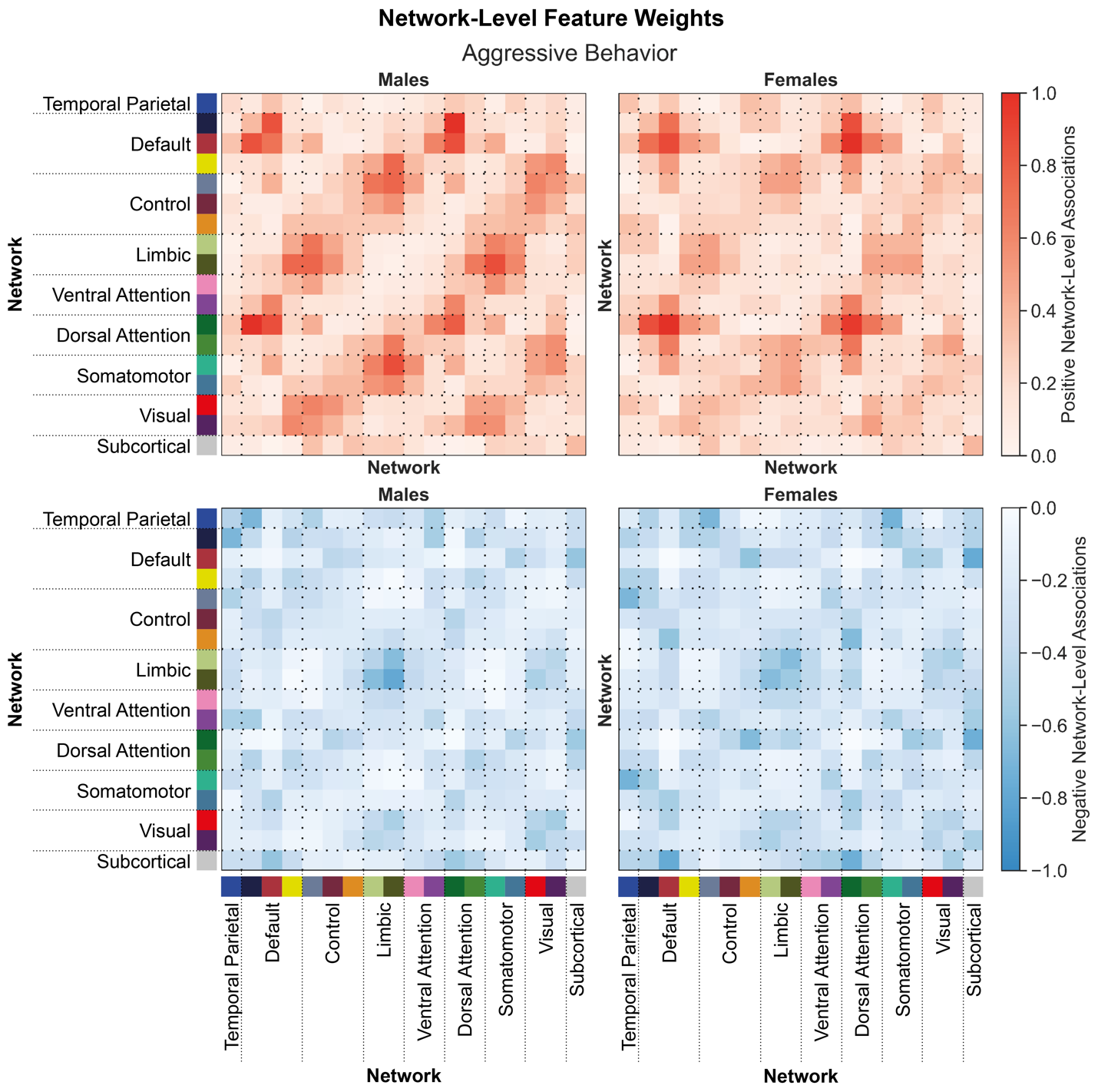
 **Supplementary Figure 5: Shared network-level functional connections underlying aggressive behaviors in males and females.**

Positive (top) and negative (bottom) associations between network-level functional connectivity and aggressive behaviors in males (left) and females (right). Regional feature weights were summarized to a network-level by assigning cortical regions to one of 17 Yeo networks, and subcortical regions to a subcortical network. Colors next to the network labels along the vertical and horizontal axes correspond to the network colors from Figure 1C. Warmer colors within the heatmap indicate a positive association and cooler colors indicate a negative association. For visualization, values within each matrix were divided by the absolute maximum value across the positive and negative matrices for each sex. Correlations between positive associations across sexes, r_positive_=0.80. Correlations between negative associations across sexes, r_negative_=0.75.

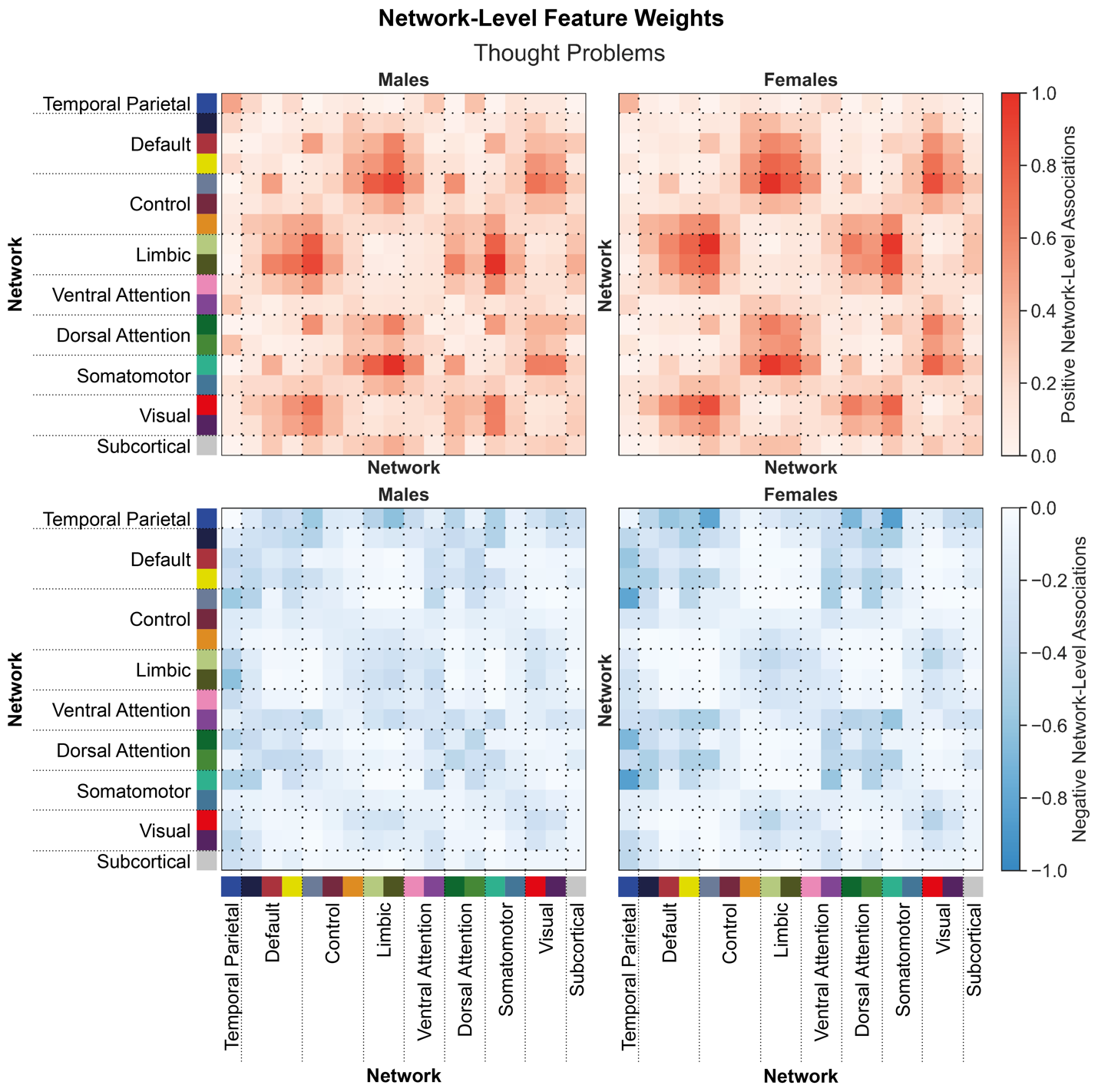
 **Supplementary Figure 6: Shared network-level functional connections underlying thought problems in males and females.**

Positive (top) and negative (bottom) associations between network-level functional connectivity and thought problems in males (left) and females (right). Regional feature weights were summarized to a network-level by assigning cortical regions to one of 17 Yeo networks, and subcortical regions to a subcortical network. Colors next to the network labels along the vertical and horizontal axes correspond to the network colors from Figure 1C. Warmer colors within the heatmap indicate a positive association and cooler colors indicate a negative association. For visualization, values within each matrix were divided by the absolute maximum value across the positive and negative matrices for each sex. Correlations between positive associations across sexes, r_positive_=0.91. Correlations between negative associations across sexes, r_negative_=0.86.

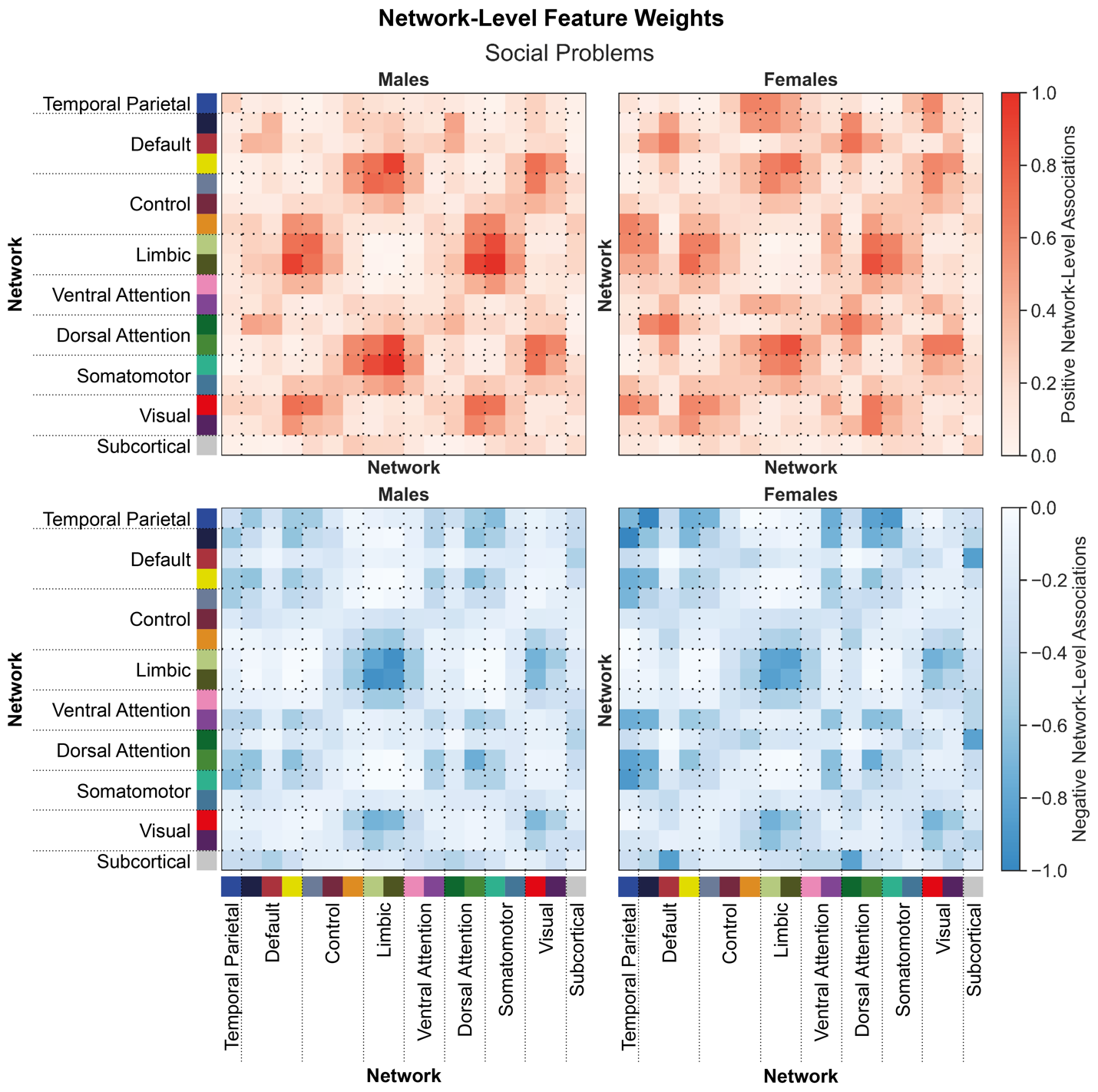
 **Supplementary Figure 7: Shared network-level functional connections underlying social problems in males and females.**

Positive (top) and negative (bottom) associations between network-level functional connectivity and social problems in males (left) and females (right). Regional feature weights were summarized to a network-level by assigning cortical regions to one of 17 Yeo networks, and subcortical regions to a subcortical network. Colors next to the network labels along the vertical and horizontal axes correspond to the network colors from Figure 1C. Warmer colors within the heatmap indicate a positive association and cooler colors indicate a negative association. For visualization, values within each matrix were divided by the absolute maximum value across the positive and negative matrices for each sex. Correlations between positive associations across sexes, r_positive_=0.82. Correlations between negative associations across sexes, r_negative_=0.90.

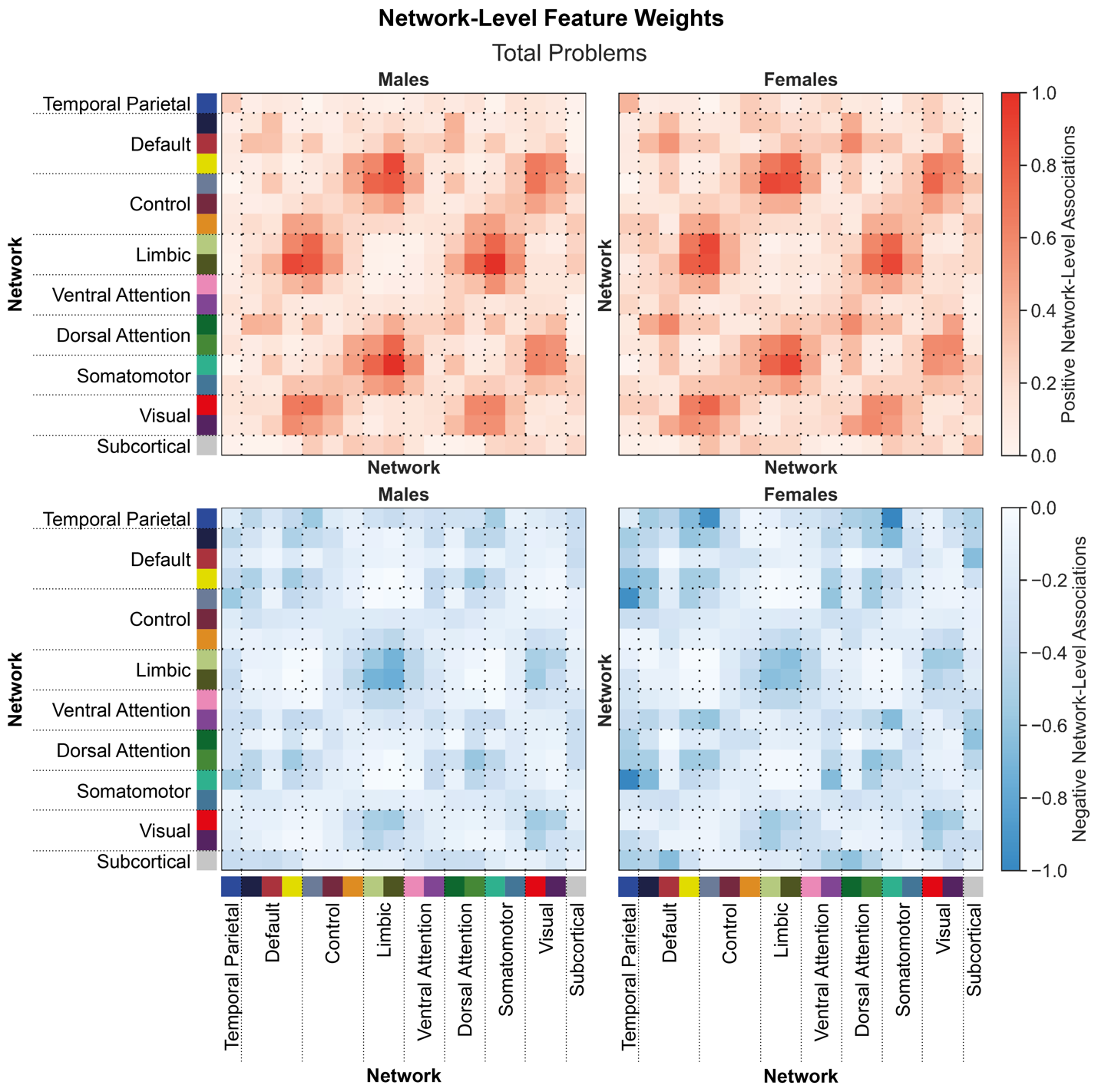
 **Supplementary Figure 8: Shared network-level functional connections underlying total problems in males and females.**

Positive (top) and negative (bottom) associations between network-level functional connectivity and total problems in males (left) and females (right). Regional feature weights were summarized to a network-level by assigning cortical regions to one of 17 Yeo networks, and subcortical regions to a subcortical network. Colors next to the network labels along the vertical and horizontal axes correspond to the network colors from Figure 1C. Warmer colors within the heatmap indicate a positive association and cooler colors indicate a negative association. For visualization, values within each matrix were divided by the absolute maximum value across the positive and negative matrices for each sex. Correlations between positive associations across sexes, r_positive_=0.95. Correlations between negative associations across sexes, r_negative_=0.88.

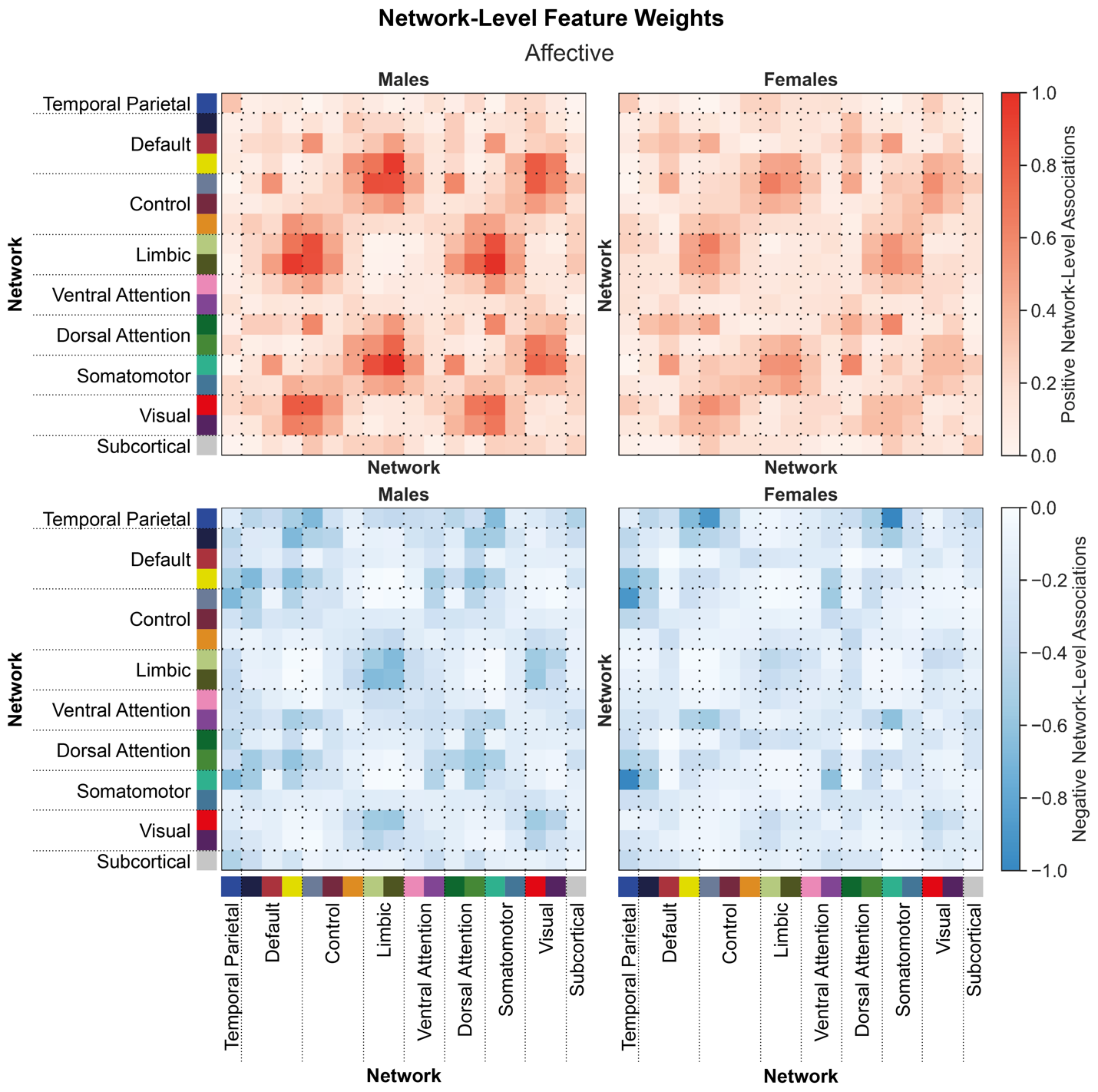
 **Supplementary Figure 9: Shared network-level functional connections underlying affective scores in males and females.**

Positive (top) and negative (bottom) associations between network-level functional connectivity and affective scores in males (left) and females (right). Regional feature weights were summarized to a network-level by assigning cortical regions to one of 17 Yeo networks, and subcortical regions to a subcortical network. Colors next to the network labels along the vertical and horizontal axes correspond to the network colors from Figure 1C. Warmer colors within the heatmap indicate a positive association and cooler colors indicate a negative association. For visualization, values within each matrix were divided by the absolute maximum value across the positive and negative matrices for each sex. Correlations between positive associations across sexes, r_positive_=0.81. Correlations between negative associations across sexes, r_negative_=0.75.

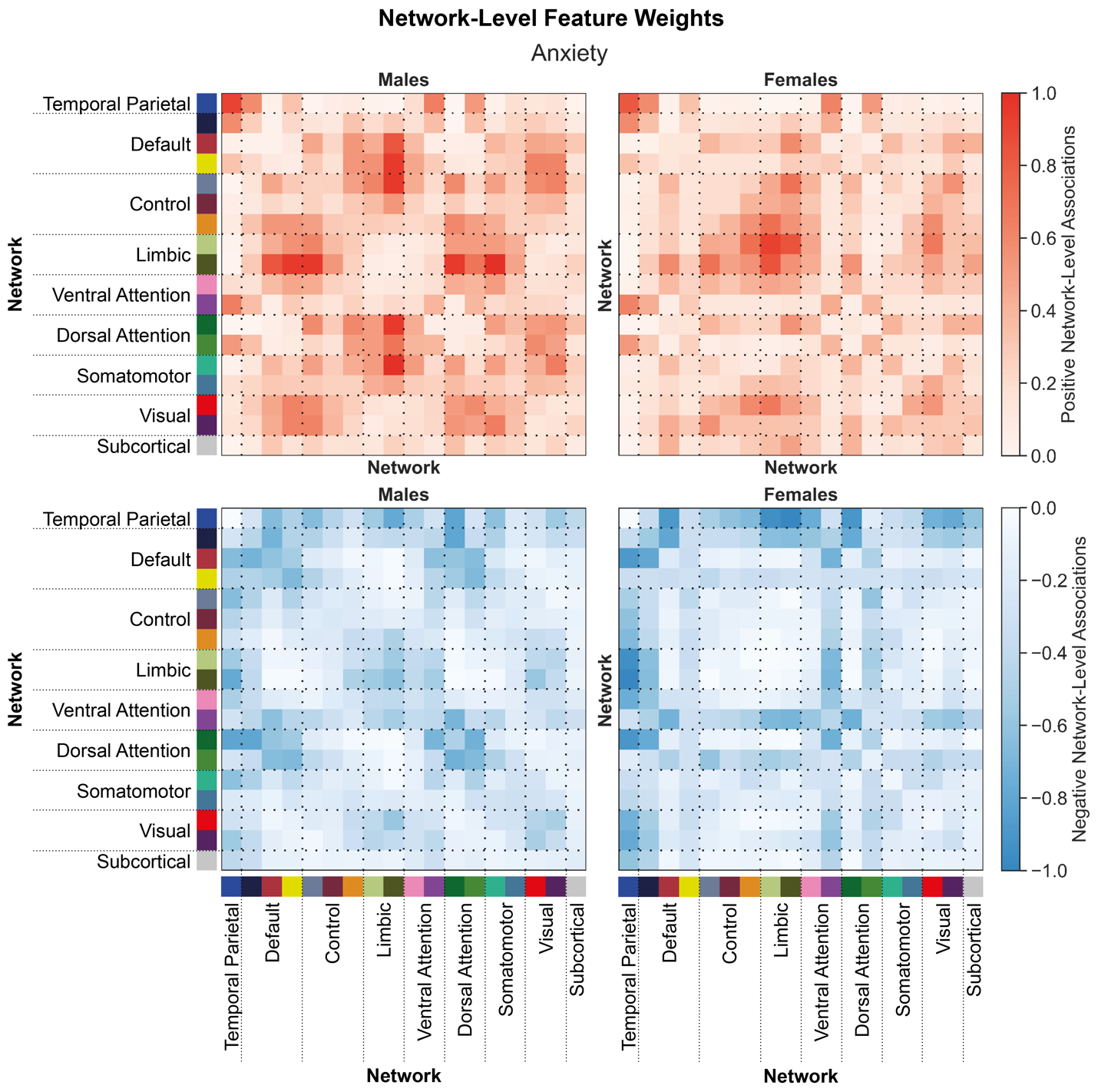
 **Supplementary Figure 10: Shared network-level functional connections underlying anxiety scores in males and females.**

Positive (top) and negative (bottom) associations between network-level functional connectivity and anxiety scores in males (left) and females (right). Regional feature weights were summarized to a network-level by assigning cortical regions to one of 17 Yeo networks, and subcortical regions to a subcortical network. Colors next to the network labels along the vertical and horizontal axes correspond to the network colors from Figure 1C. Warmer colors within the heatmap indicate a positive association and cooler colors indicate a negative association. For visualization, values within each matrix were divided by the absolute maximum value across the positive and negative matrices for each sex. Correlations between positive associations across sexes, r_positive_=0.38. Correlations between negative associations across sexes, r_negative_=0.53.

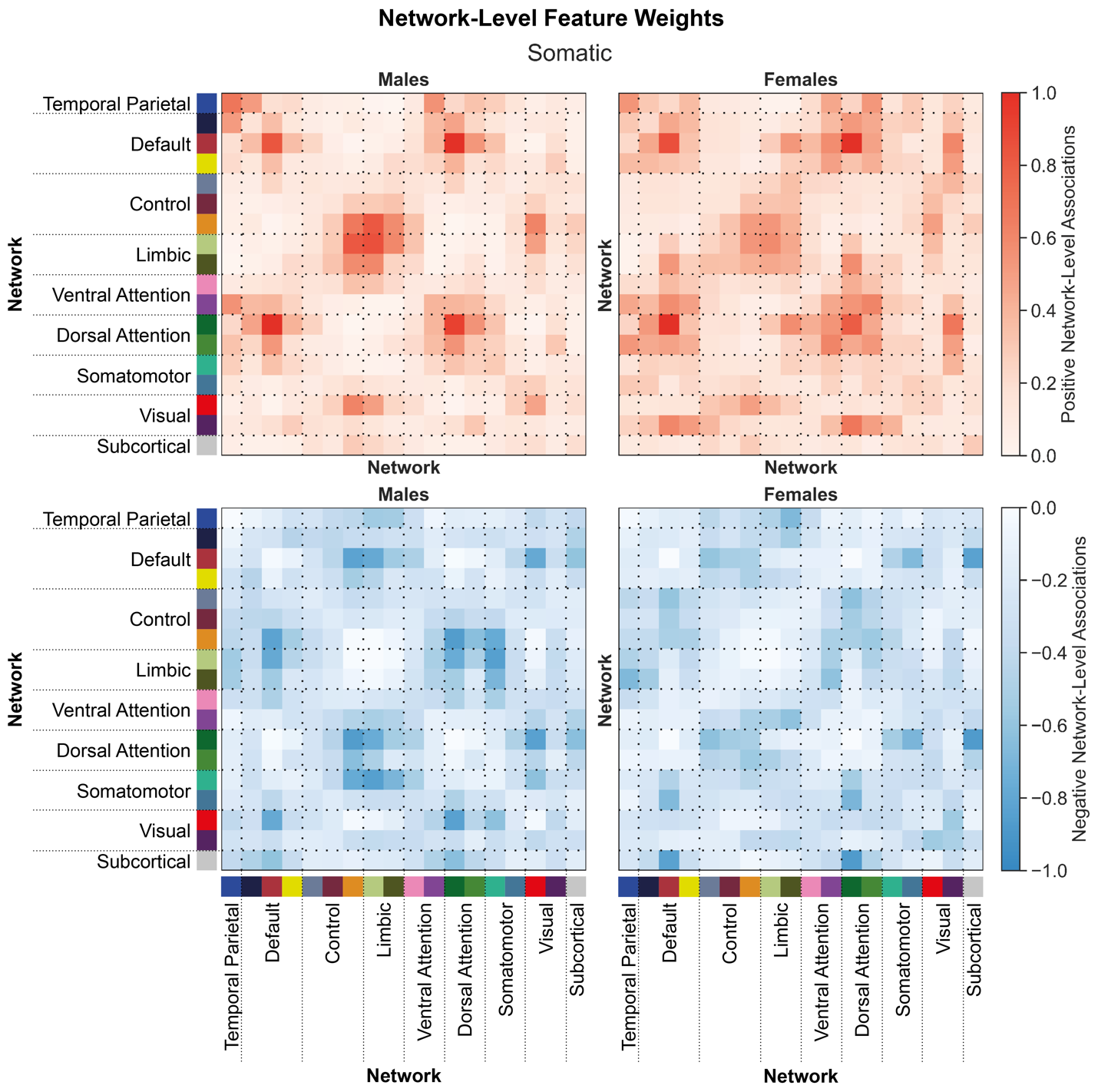
 **Supplementary Figure 11: Shared network-level functional connections underlying somatic scores in males and females.**

Positive (top) and negative (bottom) associations between network-level functional connectivity and somatic scores in males (left) and females (right). Regional feature weights were summarized to a network-level by assigning cortical regions to one of 17 Yeo networks, and subcortical regions to a subcortical network. Colors next to the network labels along the vertical and horizontal axes correspond to the network colors from Figure 1C. Warmer colors within the heatmap indicate a positive association and cooler colors indicate a negative association. For visualization, values within each matrix were divided by the absolute maximum value across the positive and negative matrices for each sex. Correlations between positive associations across sexes, r_positive_=0.76. Correlations between negative associations across sexes, r_negative_=0.56.

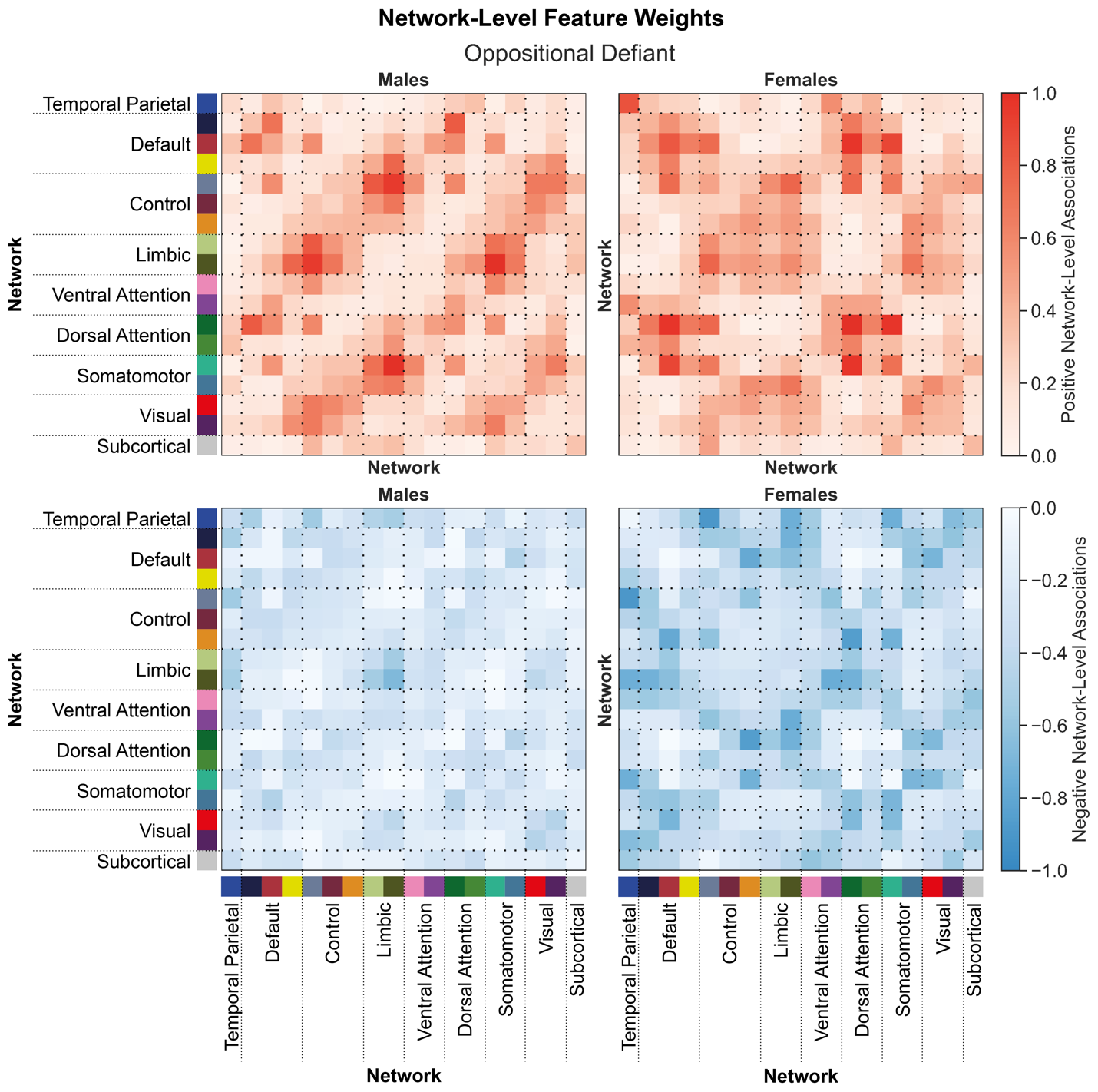
 **Supplementary Figure 12: Shared network-level functional connections underlying oppositional defiant scores in males and females.**

Positive (top) and negative (bottom) associations between network-level functional connectivity and oppositional defiant scores in males (left) and females (right). Regional feature weights were summarized to a network-level by assigning cortical regions to one of 17 Yeo networks, and subcortical regions to a subcortical network. Colors next to the network labels along the vertical and horizontal axes correspond to the network colors from Figure 1C. Warmer colors within the heatmap indicate a positive association and cooler colors indicate a negative association. For visualization, values within each matrix were divided by the absolute maximum value across the positive and negative matrices for each sex. Correlations between positive associations across sexes, r_positive_=0.50. Correlations between negative associations across sexes, r_negative_=0.27.

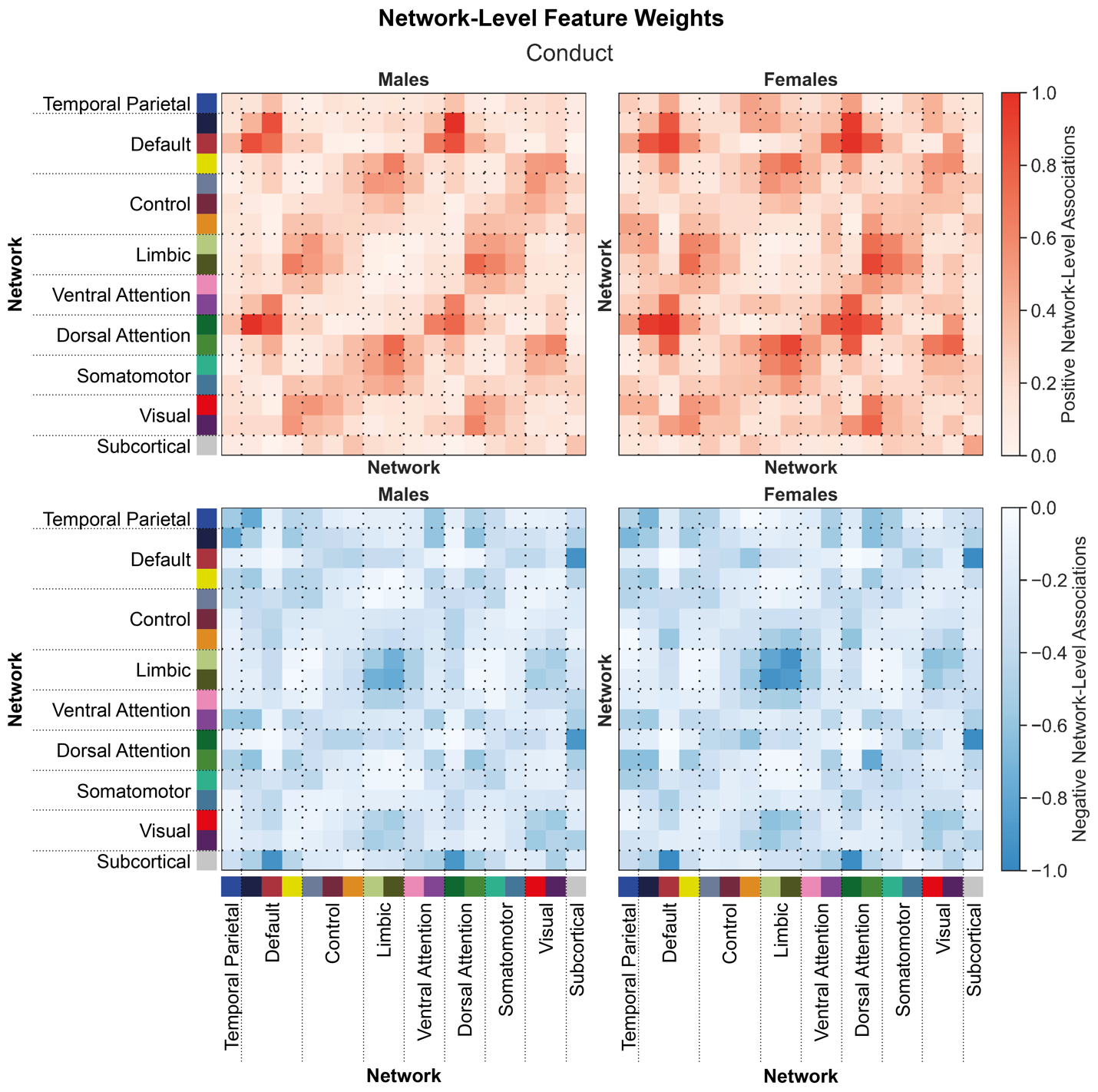
 **Supplementary Figure 13: Shared network-level functional connections underlying conduct scores in males and females.**

Positive (top) and negative (bottom) associations between network-level functional connectivity and conduct scores in males (left) and females (right). Regional feature weights were summarized to a network-level by assigning cortical regions to one of 17 Yeo networks, and subcortical regions to a subcortical network. Colors next to the network labels along the vertical and horizontal axes correspond to the network colors from Figure 1C. Warmer colors within the heatmap indicate a positive association and cooler colors indicate a negative association. For visualization, values within each matrix were divided by the absolute maximum value across the positive and negative matrices for each sex. Correlations between positive associations across sexes, r_positive_=0.89. Correlations between negative associations across sexes, r_negative_=0.93.

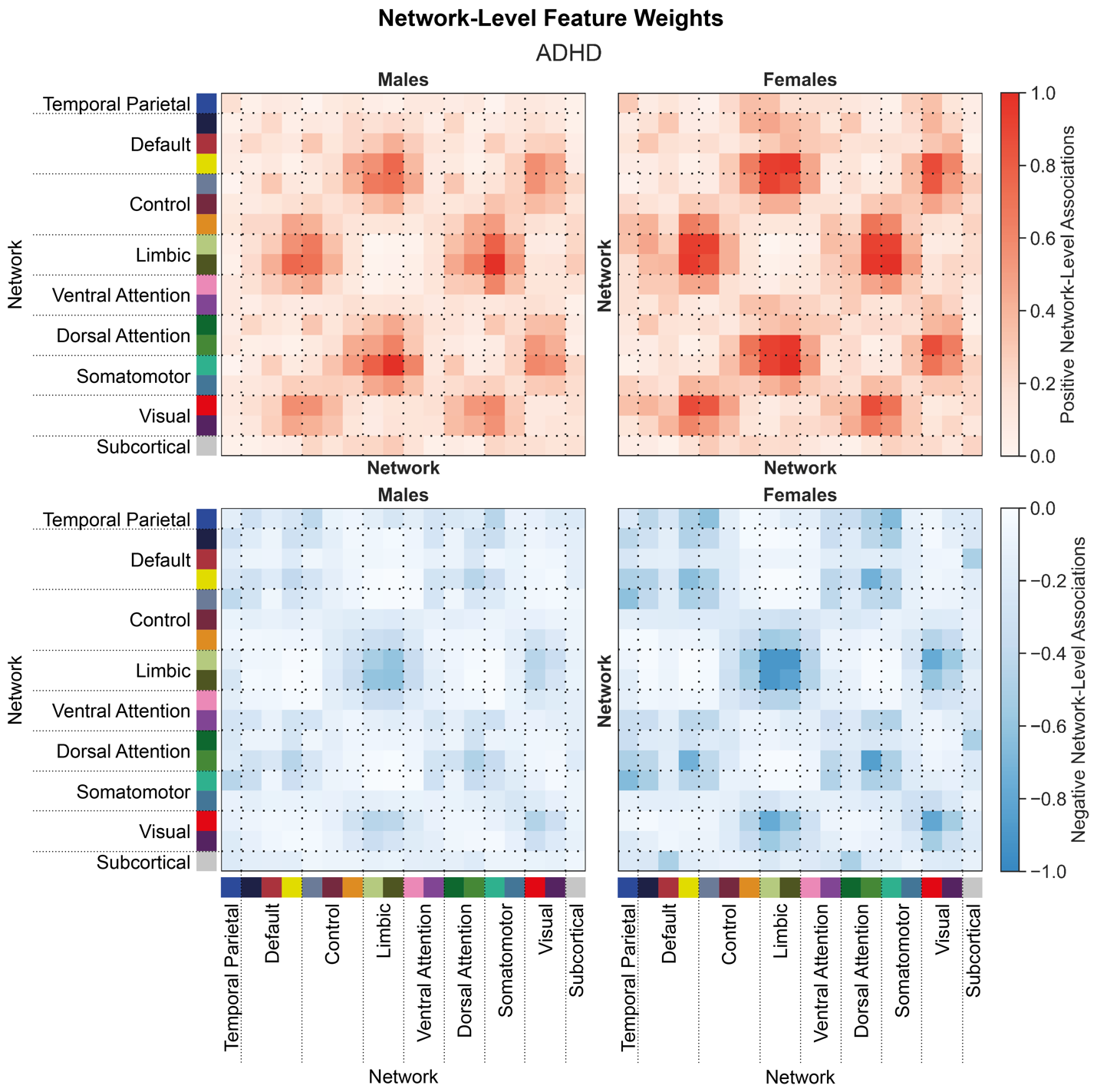
 **Supplementary Figure 14: Shared network-level functional connections underlying ADHD scores in males and females.**

Positive (top) and negative (bottom) associations between network-level functional connectivity and ADHD scores in males (left) and females (right). Regional feature weights were summarized to a network-level by assigning cortical regions to one of 17 Yeo networks, and subcortical regions to a subcortical network. Colors next to the network labels along the vertical and horizontal axes correspond to the network colors from Figure 1C. Warmer colors within the heatmap indicate a positive association and cooler colors indicate a negative association. For visualization, values within each matrix were divided by the absolute maximum value across the positive and negative matrices for each sex. Correlations between positive associations across sexes, r_positive_=0.91. Correlations between negative associations across sexes, r_negative_=0.94.
